# Single Nucleus Multiomic Profiling Reveals Age-Dynamic Regulation of Host Genes Associated with SARS-CoV-2 Infection

**DOI:** 10.1101/2020.04.12.037580

**Authors:** Allen Wang, Joshua Chiou, Olivier B Poirion, Justin Buchanan, Michael J Valdez, Jamie M Verheyden, Xiaomeng Hou, Minzhe Guo, Jacklyn M Newsome, Parul Kudtarkar, Dina A Faddah, Kai Zhang, Randee E Young, Justinn Barr, Ravi Misra, Heidie Huyck, Lisa Rogers, Cory Poole, Jeffery A. Whitsett, Gloria Pryhuber, Yan Xu, Kyle J Gaulton, Sebastian Preissl, Xin Sun, NHLBI LungMap Consortium

## Abstract

Respiratory failure is the leading cause of COVID-19 death and disproportionately impacts adults more than children. Here, we present a large-scale snATAC-seq dataset (90,980 nuclei) of the human lung, generated in parallel with snRNA-seq (46,500 nuclei), from healthy donors of ~30 weeks, ~3 years and ~30 years of age. Focusing on genes implicated in SARS-CoV-2 cell entry, we observed an increase in the proportion of alveolar epithelial cells expressing *ACE2* and *TMPRSS2* in adult compared to young lungs. Consistent with expression dynamics, 10 chromatin peaks linked to *TMPRSS2* exhibited significantly increased activity with age and harbored IRF and STAT binding sites. Furthermore, we identified 14 common sequence variants in age-increasing peaks with predicted regulatory function, including several associated with respiratory traits and *TMPRSS2* expression. Our findings reveal a plausible contributor to why children are more resistant to COVID-19 and provide an epigenomic basis for transferring this resistance to older populations.

## INTRODUCTION

Aside from fulfilling gas-exchange functions that are vital for survival beginning with the first breath, the lung functions as a critical barrier to protect against inhaled pathogens such as viruses. As the COVID-19 pandemic swept across the world, the lung came into focus because acute respiratory distress (ARDS) is the primary cause of mortality. Thus, understanding how SARS-CoV-2 infects and impacts the lung has become an urgent call-to-action.

The lung is composed of an elaborate airway tree that conducts air to and from the distal gas-exchange units called the alveoli. In an average human adult lung, an estimated 480 million alveoli give rise to approximately 1,000 ft^2^ of gas-exchange surface area (Ochs et al., 2004). Airway and alveolar epithelium constitute the respiratory barrier that is exposed to inhaled pathogens. Respiratory epithelial cells are thereby at the frontline of infection, although pathogens that have bypassed the barrier can infect other cell types. The human airway epithelium is composed of luminal cells and basal cells. Luminal cells include club cells and goblet cells that moisturize the air and trap pathogens, as well as ciliated cells that sweep out inhaled particles. These luminal cells are underlined by basal cells, which serve as progenitors when luminal cells are lost after infection. The alveolar epithelium is composed of alveolar type 1 cells (AT1s) which line the gas-blood interface and alveolar type 2 cells (AT2s) which produce surfactant to reduce surface tension and protect against pathogens. While SARS-CoV-2 likely infects both the airway and alveolar regions of the lung, it is the damage to the alveolar region that underlines acute respiratory distress syndrome (Du et al., 2020).

Several large scale studies including efforts from LungMap and the Human Cell Atlas aim to generate a map of cell types in the human lung with single cell transcriptomics as the central modality (Reyfman et al., 2019; Schiller et al., 2019; Travaglini et al., 2020; Xu et al., 2016). Regions of the human genome, such as promoters or distal enhancers, can regulate cell-type specific gene expression in *cis* (Consortium, 2012; Roadmap Epigenomics et al., 2015; Thurman et al., 2012). Accessible or ‘open’ chromatin is a hallmark of *cis*-regulatory elements, and can be assayed using techniques such as DNase-seq and ATAC-seq (Buenrostro et al., 2013; Thurman et al., 2012). To overcome tissue heterogeneity single cell technologies like single cell ATAC-seq have been developed to map the epigenome and gene regulatory programs in component cell types within heterogeneous tissues (Buenrostro et al., 2015; Chen et al., 2018; Cusanovich et al., 2015; Cusanovich et al., 2018; Lareau et al., 2019; Satpathy et al., 2019). Profiles derived from single cells can elucidate cell type-specific *cis*-regulatory elements, transcriptional regulators driving element activity, and predicted target genes of distal elements using single cell co-accessibility (Cusanovich et al., 2018; Lareau et al., 2019; Pliner et al., 2018; Preissl et al., 2018; Satpathy et al., 2019). Human sequence variants affecting susceptibility to complex physiological and disease traits are enriched in non-coding sequence (Maurano et al., 2015; Pickrell, 2014), and cell type-specific profiles derived from single cell epigenomic data can help prioritize cell types of action for these variants (Chiou et al., 2019; Corces et al., 2020).

Both *in silico* structural modeling as well as biochemical assays have implicated several key host proteins at the top of the hierarchy for SARS-CoV-2 infection. ACE2 has been demonstrated as the receptor for not only the original SARS-CoV, but also SARS-CoV-2 (Lan et al., 2020; Yan et al., 2020). Based mainly on literature from the original SARS-CoV as well as emerging data from SARS-CoV-2 (Huang et al., 2006; Matsuyama et al., 2020; Reinke et al., 2017; Walls et al., 2020; Zhou et al., 2016), TMPRSS2 and CTSL are responsible for fusion of the virus with host cell by cleaving the viral Spike protein. BSG is a receptor that can bind to the SARS-CoV spike protein (Chen et al., 2005) and SARS-CoV-2 contains a novel cleavage site for the protease Furin, adding both genes to the list of host machinery highjacked by the virus (Coutard et al., 2020; Walls et al., 2020). In this study, we will focus on the genes encoding these 5 proteins, *ACE2, TMPRSS2, CTSL, BSG*, and *FURIN*, and determine their expression and associated epigenomic landscape at single cell resolution in the non-diseased human lung.

In the race to control the COVID-19 pandemic, there has been a tremendous collective effort from the research community to elucidate the mechanism underlying SARS-CoV-2 infection. Our study contributes to this effort through a unique dataset profiling the human lung. First, we generated single cell data across neonatal, pediatric, and adult lungs from three donors in each group. These data allowed us to assess age-associated changes with minimal technical variation. Second, from each lung sample, we generated parallel snRNA-seq and snATAC-seq data. This combination allowed us to associate cell type-specific accessible chromatin profiles that may act as *cis*-regulatory regions that control cell-type specific gene expression. Using these data, we first addressed cell-type specificity and temporal dynamics of *ACE2, TMPRSS2, CTSL, BSG*, and *FURIN* expression. We next identified candidate *cis*-regulatory elements co-accessible with the promoters of these genes and characterized their cell-type specificity and temporal dynamics. Finally, we profiled sequence variation that may impact *cis*-regulatory element activity and contribute to differential susceptibility to SARS-CoV-2 infection.

Emerging epidemiology data, including on US cases reported by the CDC, demonstrate that many fewer children tested positive for SARS-CoV-2 infection, and those who tested positive generally show less severe symptoms than adults or elderly individuals (Bi et al., 2020; CDC, 2020). This age divide coincides with the finding that normal lung development in humans continues until the early 20s (Narayanan et al., 2012). Therefore COVID-19 preferentially impacts fully mature lungs relative to developing lungs. Widespread speculation has attempted to explain these age-associated differences, including immune senescence in the aging population. Defining the mechanism underlying the apparent resistance of children to COVID-19 will inform how we can transfer this resistance to adult and elderly populations.

## RESULTS

### Single nucleus RNA-seq and ATAC-seq data generation

To profile cell type specific gene expression and accessible chromatin dynamics in the human lung, we performed single nucleus RNA-seq (snRNA-seq) and single nucleus ATAC-seq (snATAC-seq) of non-diseased human lung tissue from donors of three age groups: ~30 week old gestational age (GA, prematurely born, 30wk^GA^), ~3 year old (3yo), and ~30 year old (30yo) (Supplementary Table 1). Three lungs were sampled for each age group, with both males and females represented (Supplementary Table 1). Of the 9 donors, 5 were Caucasian, 1 was African American and 3 were of unknown ancestry. For all samples, flash frozen biopsies from equivalent small airway regions of the lung were used. Nuclei were isolated from individual biopsies and split into two pools, one for snRNA-seq and one for snATAC-seq. For snATAC-seq, we generated technical replicates for one of the 3yo donors (D032) and an additional dataset for a lung sample from a 4-month-old donor (Supplementary Table 1).

To generate snRNA-seq libraries, we used the droplet-based Chromium Single Cell 3’ solution (10x Genomics) (Zheng et al., 2017). The datasets showed a clear separation of nuclei from background in the knee plot (Figure S1A). The average number of nuclei that passed initial quality control filtering per sample was 6,676 for 30wk^GA^, 7,379 for 3yo, and 4,217 for 30yo (Figure S1B). Since we profiled nuclei with a high fraction of nascent, unspliced RNA molecules, sequencing reads were mapped to an exon+intron reference. We detected on average 1,662 gene/nuclei for 30wk^GA^, 1,394 for 3yo and 1,260 for 30yo (Figure S1C). Libraries were sequenced to comparable saturation (58.4 % for 30wk^GA^, 51.6 % for 3yo and 55.0 % for 30yo; Fig. S1D).

For snATAC-seq library generation we used a semi-automated combinatorial barcoding platform (Cusanovich et al., 2015; Fang et al., 2019; Preissl et al., 2018). For each dataset, nuclei with >1,000 uniquely mapped sequencing reads were included in the analysis (Fig. S1E). The average number of nuclei that passed this threshold per age group was 8,691 for 30wk^GA^, 7,877 for 3yo and 8,034 for 30yo (Fig. S1F). The average number of reads per nucleus was 6,399 for 30wk^GA^, 7,199 for 3yo and 8,362 for 30yo (Fig. S1G). The fraction of reads in peaks (FRiP) on average per data set was 52.8 % for 30wk^GA^, 54.4 % for 3yo and 45.6 % for 30yo (Fig. S1H). These values indicate consistently high signal to noise ratios for all libraries.

### Age-linked increase in host genes for SARS-CoV-2 entry

In total, 46,500 single nucleus transcriptomes were included in the analysis after filtering out low quality nuclei and potential barcode collisions (Figure S1, Supplementary Table 2, see **Methods**). Following batch correction all datasets were merged, and 31 clusters were identified (Figure 1A). These clusters represented all major cell types in the small airway region of the lung, as well as rare cell types such as pulmonary neuroendocrine cells (Figure 1A, Figure S2A, Supplementary Table 2). We identified 14,527 epithelial cells (31.2 % of all nuclei) in our snRNA-seq dataset. This optimal representation and large number of cells allowed us power to profile gene expression patterns of viral entry genes in lung epithelial cells. For downstream analysis, we excluded an unclassified cluster, and enucleated erythrocytes because the latter were only detected in a single neonatal sample, consistent with immaturity (Figure 1A).

**Figure 1.**
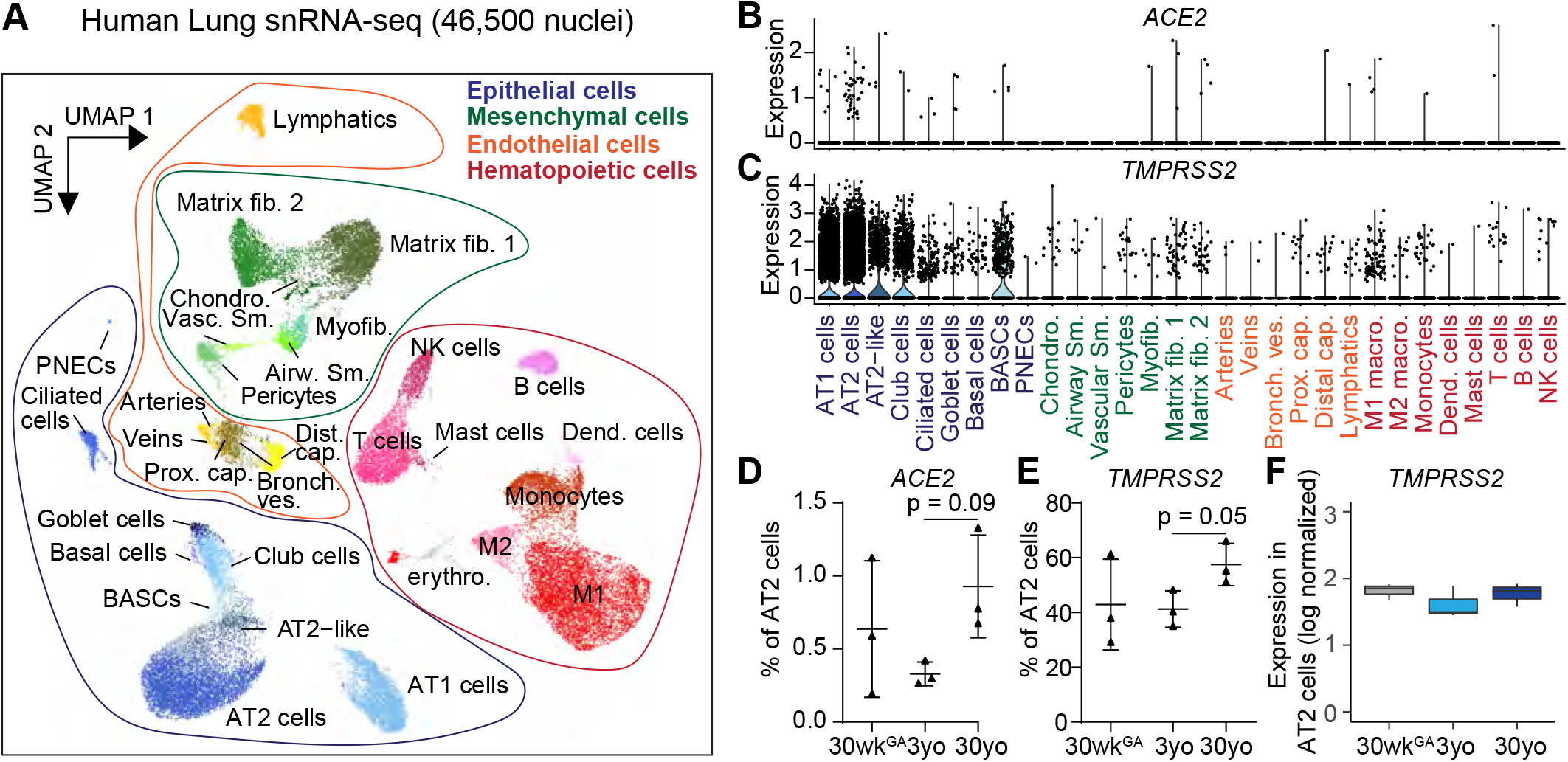
snRNA-seq of human lungs reveals expression of SARS-CoV-2 cell entry genes in the epithelial cell lineage. **A** UMAP embedding and clustering result of 46,500 snRNA-seq data from 9 donors (Premature born (30 week^GA^ of pregnancy), 3 yo, 30 yo; n = 3 per time point) identifies 31 clusters. Each dot represents a nucleus. Spread-out grey dots correspond to nuclei of unclassified cluster. **B, C** Cluster specific violin blots of gene expression of **B** *ACE2* and **C** *TMPRSS2*. **D, E** Fraction of AT2 cells with expression of *ACE2* and *TMPRSS2* at each time point. All data are represented as mean ± SD. p values derived from t-tests; One-way ANOVA did not reach significance. **F** Box plot of log normalized expression of *TMPRSS2* in AT2 cells at each time point. Displayed are the median expression values for AT2 nuclei in individual samples with at least 1 UMI.

Focusing on SARS-CoV-2 viral entry genes, we found that *ACE2* transcript was detected in very few nuclei (total 80 nuclei) in the normal lung and these nuclei were enriched within the epithelial lineage (Figure 1B, Supplementary Table 2). Alveolar type 2 (AT2) cells had the highest number of *ACE2*^+^ nuclei, accounting for 48.8% of all *ACE2*-expressing nuclei (39 out of total 80 *ACE2*^+^ nuclei). In comparison, *TMPRSS2* transcript was detected more frequently (e.g. in 3,315/7,226 nuclei, or 45.8% of the AT2 cells, Figure 1C, Supplementary Table 2). Most *TMPRSS2*-expressing cells were epithelial cells including alveolar type 1 and 2 (AT1, AT2) cells and airway cells such as club, ciliated and goblet cells (Figure 1C, Supplementary Table 2). We also detected significant correlation between the fraction of *ACE2*^+^ and *TMPRSS2*^+^ AT2 nuclei (Figure S2E) and found 21 of the 39 ACE^+^ AT2 cells also expressed *TMPRSS2* (Supplementary Table 2). The other three candidate genes of SARS-CoV-2 host cell entry *CTSL, BSG* and *FURIN* were expressed in a large number of AT1, AT2, matrix fibroblast, and M1 macrophage cells, as well as a small number of additional cell types (Figure S2B-D, Supplementary Table 2). These findings suggest that among cells that constitute the barrier exposed to inhaled pathogens, cell types in both the airway and alveolar epithelium express genes critical for SARS-CoV-2 entry.

We next asked if there were genes enriched in *ACE2*^+^ AT2 cells as compared to *ACE2*^-^ AT2 cells to identify potentially co-expressed genes. Among genes that showed a trend for higher expression in *ACE2*^+^ compared to *ACE2*^-^ cells was *IFNGR1* (log2 (fold change) = 0.4, −log_10_(p-value)=5.0; FDR corrected p=0.257, Supplementary Table 3), raising the possibility that *ACE2* may be co-regulated with interferon pathway genes, in line with conclusions of a recent study (Ziegler, 2020). In our data generated from normal lungs this correlation was modest, suggesting there is low baseline co-expression of *ACE2* and *IFNGR1*. Among genes with increased expression in *TMPRSS2*^+^ versus *TMPRSS2*^-^ AT2 cells was *ICAM1* (log2 (fold change)=0.27, −log_10_(FDR corrected p)=12.2, Supplementary Table 3), which encodes a receptor for Rhinovirus (Zhou et al., 2017). The potential co-expression of *TMPRSS2* and *ICAM1* may contribute to the often-observed co-infection by more than one respiratory virus. Indeed, co-infection of SARS-CoV-2 and other viruses including Rhinovirus has been observed, promoting urgent calls to halt the clinical practice of using positive test for other respiratory viruses as an indicator for the absence of coronavirus infection (Wang et al., 2020; Wu et al., 2020). To gain additional insight into the potential mechanisms of co-infection, we interrogated the expression of a number of known factors, receptors and proteases that have been implicated in viral entry for several key respiratory viruses (Figure S3)(Battles and McLellan, 2019; Bochkov and Gern, 2016; Laporte and Naesens, 2017; Peck et al., 2015). For examples, consistent with prior findings, we found that *CDHR3*, a receptor for Rhinovirus C, was expressed most abundantly in ciliated cells (Battles and McLellan, 2019; Bochkov and Gern, 2016; Laporte and Naesens, 2017; Peck et al., 2015). *ANPEP*, the entry receptor for HCoV-229E, was predominantly expressed in macrophages and to a lesser extent in club and other epithelial cells (Waradon Sungnak, 2020; Yeager et al., 1992). Compared to *ACE2, DPP4*, which encodes the host receptor for MERS-CoV, was detected much more frequently overall, and especially in AT2, AT1 and T cells (Figure S3) (Raj et al., 2013; Waradon Sungnak, 2020). This single cell resolved view may contribute to a comprehensive map of the routes of respiratory viral entry.

The leading cause of death for COVID-19 is Acute Respiratory Distress Syndrome (ARDS) which is characterized by failure of gas-exchange due to destruction of the alveolar region of the lung (Du et al., 2020). AT2 is an abundant epithelial cell type in the alveolar region and expresses all of the SARS-CoV-2 viral entry genes assayed here and likely bears the brunt of infection. Consequently, we focused on AT2 cells for follow up analysis. We found that the percentage of AT2 cells expressing *ACE2* had an increasing trend in 30yo adult samples compared to 3yo samples (Figure 1D). In addition, we found a strong trend of increase in the percentage of AT2 cells expressing *TMPRSS2* in adult samples compared to 3yo samples (41.2 ± 6.6% for 3yo and 57.4 ± 7.7% for 30yo, p = 0.05 (t-test), Figure 1E). While very few *ACE2^+^/TMPRSS2*^+^ double positive AT2 nuclei were detected, the fraction of these nuclei in all AT2s increased with age (0.2 % (6 nuclei) in 30wk^GA^, 0.3% (5 nuclei) in 3yo and 0.5% (10 nuclei) in 30yo, Supplementary Table 2). Of note, one of the samples in the 30wk^GA^ cohort D062 appeared to be an outlier in its expression of multiple analyzed genes. A review of pathology notes revealed mild features of respiratory distress syndrome including epithelial autolysis and increased alveolar macrophages in this sample, suggesting potential reasons for the variation. In a supplementary analysis, excluding this sample resulted in stronger age-associated effects (Figure S2F, G). For example, there was a significant increase in the fraction of *TMPRSS2*^+^ AT2 cells between 30wk^GA^ and 30yo samples (Figure S2G).

The increase in proportion of AT2 cells expressing *ACE2* and *TMPRSS2* is unlikely due to differences in genes captured per nucleus as the adult samples had the lowest numbers of genes/nucleus, suggesting that the extent of expression increase is likely a conservative estimation (Figure S1C). In contrast to the percentage of AT2 nuclei expressing these genes, the expression levels per nucleus were not different across different age groups for either *ACE2* (no nucleus had >1 UMI detected) or *TMPRSS2* (Figure 1F). Together, an increased proportion of host cells expressed *TMPRSS2* and *ACE2* in adults, the latter just a trend due to the sparsity of *ACE2*^+^ cells, suggesting that a higher percentage of cells in the adult lung can be infected by SARS-CoV-2.

Since a large proportion of COVID-19 patients are elderly, we sought to compare viral entry gene expression in aged lungs to expression in our samples. The LungMap Human Tissue Core, which provided the frozen biopsies for this study, does not have donors older than ~30. We therefore instead, identified 4 publicly available scRNA-seq datasets from non-diseased lungs of ages >55 that served as controls in pulmonary fibrosis studies (Morse et al., 2019; Reyfman et al., 2019). We integrated snRNA-seq data from our study (n=9) with these 4 scRNA-seq samples (Supplementary Table 1) using Seurat 3 (Stuart et al., 2019). AT2 cells clustered together across all samples with minimal evidence for batch effects (Figure S4A). Compared to 30yo samples, we observed a trend for increased frequency of *ACE2*^+^ (p = 0.095) and *TMPRSS2*^+^ (p = 0.070) AT2 cells in the >55yo group (Aged; Figure S4B). While these patterns are consistent with epidemiological findings that elderly are at highest risk, we make these observations cautiously due to the multiple potential confounding variables present when comparing across independent datasets spanning multiple methodologies.

### Annotation of *cis*-regulatory sequences linked to SARS-CoV-2 viral entry gene activity

To investigate *cis*-regulatory elements driving cell-type specific and age-related patterns of SARS-CoV-2 viral entry gene expression, we examined snATAC-seq data generated from the same nuclei preparations. After batch correction and filtering of low-quality nuclei and likely doublets, we clustered and analyzed a total of 90,980 single nucleus accessible chromatin profiles. We identified 19 clusters representing epithelial (AT2, AT2, club, ciliated, basal and neuroendocrine), mesenchymal (myofibroblast, pericyte, matrix fibroblast 1 and matrix fibroblast 2), endothelial (arterial, lymphatic, and 2 clusters of capillaries), and hematopoietic cell types (macrophage, B-cell, T-cell, NK cell and enucleated erythrocyte) (Figure 2A). Supporting these cluster annotations, we observed cell type-specific patterns of chromatin accessibility at known marker genes for each cell type (Figure S5A).

**Figure 2.**
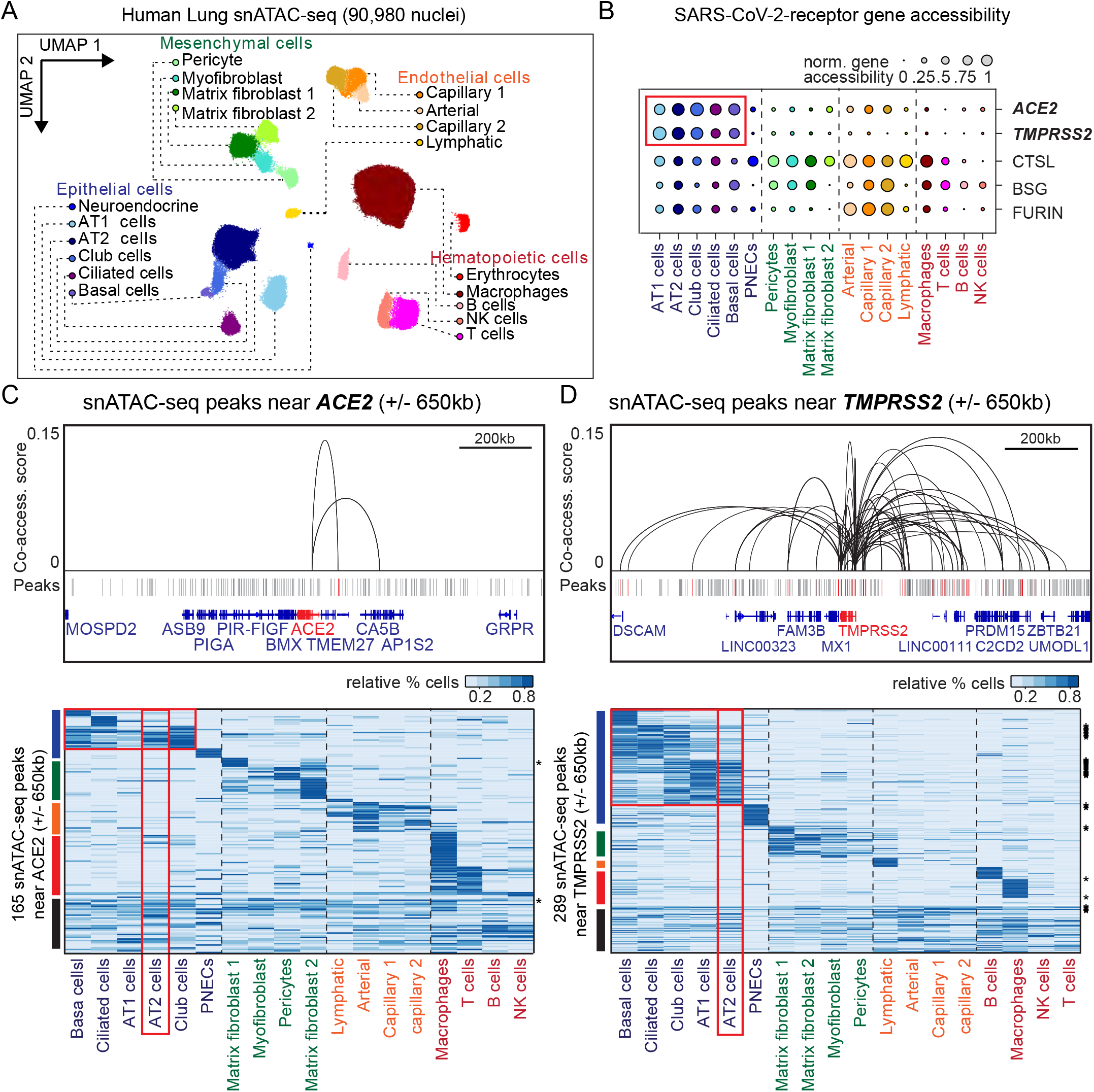
snATAC-seq analysis of human lungs reveals candidate *cis* regulatory elements for *ACE2* and *TMPRSS2*. **A** UMAP embedding and clustering results of snATAC-seq data from 90,980 single-nucleus chromatin profiles from ten donors (Premature born (30 week^GA^, n = 3), 4 month old (n = 1), 3 yo (n = 3) and 30 yo (n = 3)). **B** Gene accessibility of candidate SARS-CoV-2 cell entry genes. **C** Union set of peaks identified in all clusters surrounding *ACE2* (+/- 650 kb) and elements that show co-accessibility (co-accessibility score > 0.05) with the *ACE2* promoter via Cicero (Cusanovich et al., 2018) (top panel). Hierarchical clustering of the relative proportion of cells (see methods) with a fragment within 165 peak regions surrounding *ACE2* (lower panel). Asterisks highlight peaks co-accessible with the *ACE2* promoter via Cicero. Horizontal red box highlights peaks with increased relative accessibility shared in basal, ciliated, AT1, AT2 and club cells as compared to other cell types. Vertical red box highlights peaks with increased relative accessibility in AT2 cells. **D** Union set of peaks identified in all clusters surrounding *TMPRSS2* (+/- 650 kb) and elements that show co-accessibility with the *TMPRSS2* promoter (top panel; co-accessibility score >0.05 (Cusanovich et al., 2018)). Hierarchical clustering of the relative proportion of cells with a fragment within 289 peak regions surrounding *TMPRSS2* (lower panel). Horizontal red box highlights peaks with increased relative accessible cells shared in basal, ciliated, club, AT1 and AT2 cells as compared to other cell types. Vertical red box highlights peaks with increased relative accessibility in AT2 cells. Asterisks highlight peaks co-accessible with the *TMPRSS2* promoter.

Focusing on SARS-Cov-2 viral entry genes, both *ACE2* and *TMPRSS2* were primarily accessible throughout their gene body in alveolar cells such as AT1, AT2, and airway cells such as club, ciliated, and basal cells (Figure 2B). Conversely, the *CTSL* gene body exhibited chromatin accessibility across epithelial cells, mesenchymal cells, endothelial, and macrophages. *BSG* and *FURIN* also showed broad chromatin accessibility patterns with the highest activity in endothelial cells, such as capillaries (Figure 2B). Overall, the patterns of chromatin accessibility across cell types at genes involved in SARS-CoV-2 cell entry substantiate our conclusions from snRNA-seq data, including the finding that *ACE2* and *TMPRSS2* are primarily expressed in alveolar and airway cells (Figure 1B,C).

To identify specific *cis*-regulatory elements that might control cell type-restricted expression of the SARS-CoV-2 viral entry genes in the lung, we aggregated cells within each cell type and called accessible chromatin sites from the aggregated profiles using MACS2 (Zhang et al., 2008). We then identified sites mapping within 650kb of each SARS-CoV-2 viral entry gene, and further identified sites that were co-accessible with the gene promoter using Cicero (Pliner et al., 2018). At the *ACE2* locus, we identified 165 accessible chromatin sites mapping within the ±650kb window (Figure 2C, Supplementary Table 4). Of these 165 sites, only two were co-accessible with the *ACE2* promoter (Figure 2C, Supplementary Table 5). We speculate that the low number of co-accessible sites is likely due to the small percentage of *ACE2*^+^ nuclei (Figure 1B). When examining the accessibility of the 165 peaks at the *ACE2* locus across cell types, we observed clear sub-groupings of sites, including those specific to basal cells, specific to ciliated cells, and shared across basal, ciliated, AT1, AT2, and club cells (Figure 2C, Supplementary Table 5).

At the *TMPRSS2* locus, we identified 289 accessible chromatin sites mapping in the ±650kb window, of which 37 were co-accessible with the *TMPRSS2* promoter (Figure 2D, Supplementary Tables 4 and 5). In agreement with *TMPRSS2* gene accessibility in alveolar and airway cells, 113 out of the 289 elements exhibited patterns of accessibility specific to basal, ciliated, club, AT1, and AT2 cells. We observed a basal cell-specific cluster and two broader epithelial cell clusters (basal, ciliated, and club enriched; and club, AT1, and AT2 enriched) (Figure 2D, Supplementary Table 5). Notably, the majority of sites co-accessible with *TMPRSS2* (25/37) were found within these broad alveolar- and airway-enriched clusters suggesting that these elements are likely responsible for alveolar and airway expression of *TMPRSS2*.

Finally, at the *CTSL, FURIN*, and *BSG* loci we identified 262, 293, and 272 accessible chromatin sites, respectively, within a ±650kb window of which 6, 56, and 47 were co-accessible with their respective gene promoters (Figure S5B, C, D, Supplementary Tables 4 and 5). Sites for all three genes exhibited broad patterns of accessible chromatin signal across cell types consistent with broad accessibility across gene bodies. This collection of cell-type resolved candidate *cis*-regulatory elements associated with SARS-CoV-2 host genes will be critically important for follow up studies to determine how host cell genes are regulated and how genetic variation within these elements contributes to infection rate and disease outcomes.

### *Cis*-regulatory elements linked to *TPMRSS2* are part of an age-related regulatory program associated with immune signaling in AT2 cells

Having observed increasing percentages of *TMPRSS2* expressing cells with age in AT2 cells (Figure 1E, Figure S2G), we speculated that *TMPRSS2* may be under the control of an age-related *cis* regulatory program. To investigate whether an age-associated *cis*-regulatory network exists in AT2 cells, we identified accessible chromatin sites in AT2 cells that show dynamic accessibility across donor age groups. Based on our findings from snRNA-seq we speculate that these dynamics will be at least in part due to a higher number of cells expressing these genes rather than more activity within a cell. We tested all possible pairwise age comparisons between AT2 signal from each of the three groups of 30wk^GA^, 3yo, and 30yo donors while accounting for donor to donor variability (Figure 3A). Overall, we identified 22,745 age-linked sites in AT2 cells which exhibited significant differences (FDR<0.05) in any pairwise comparison (Figure 3A, B). Clustering of these dynamic peaks revealed five predominant groups of age-dependent chromatin accessibility patterns (cI-cV, Fig 3B).

**Figure 3.**
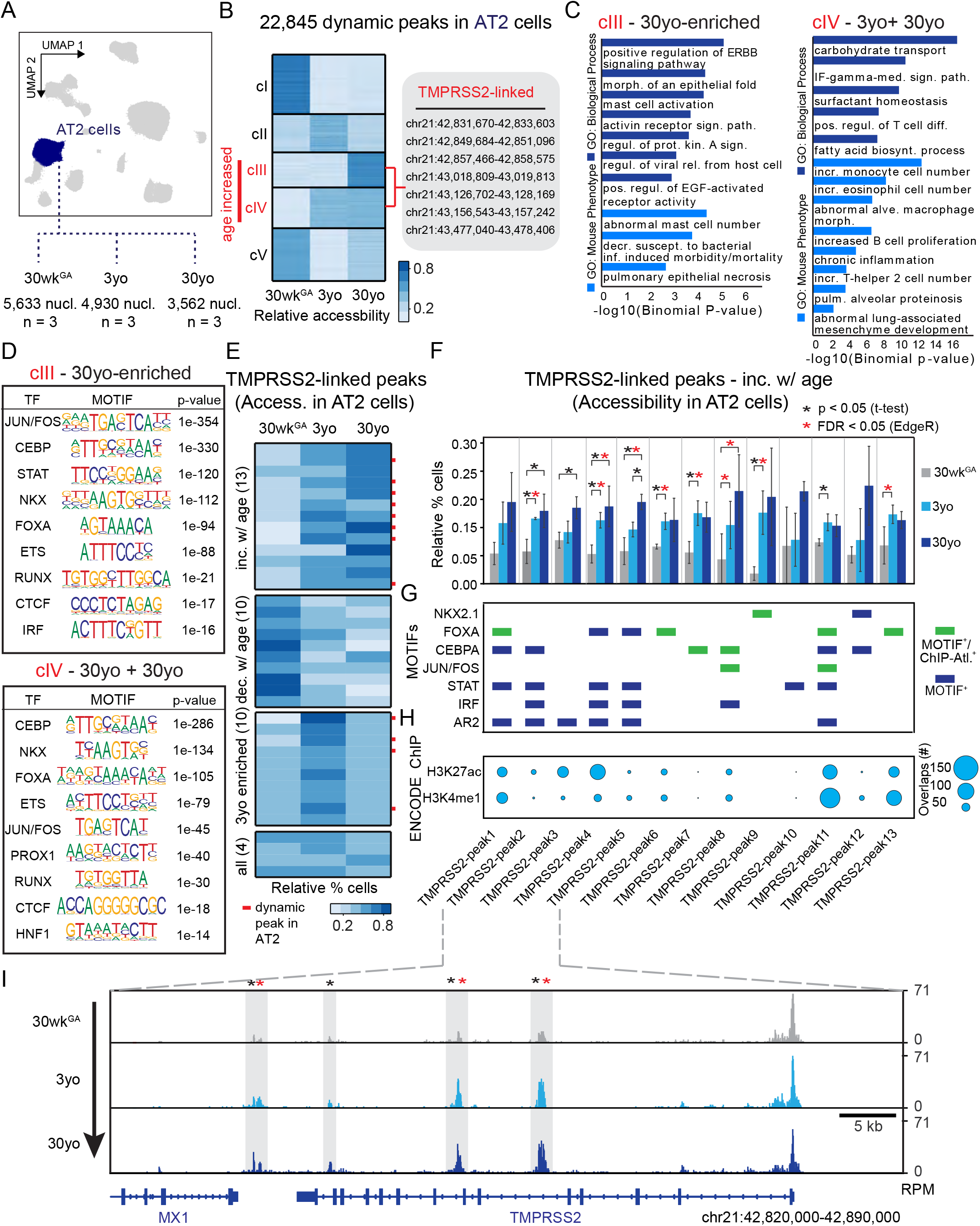
Age-increasing accessible chromatin in AT2 cells exhibits signatures of immune regulation and harbors *TMPRSS2*-linked sites of chromatin accessibility. **A** Differential analysis was performed on AT2 cells using pairwise comparisons between three ages with replicates (n = 3 per stage). **B** K-means cluster analysis (K=5) of relative accessibility scores (see **Methods**) for 22,845 age-dynamic peaks (FDR < 0.05, EdgeR) in AT2 cells. Clusters III and IV show increasing accessibility with age and contain seven *TMPRSS2*-co-accessible sites. **C** GREAT (McLean et al., 2010) analysis of elements in group cIII (left panel) and cIV (right panel) shows enrichment of immune related gene ontology terms. **D** Transcription factor motif enrichment analysis of elements in cIII and cIV. **E** Classification of age-dynamic patterns across the 37 *TMPRSS2*-co-acessible sites based on the relative percentage of AT2 cells with at least one fragment overlapping each peak. Red bars indicate dynamic peaks identified from analysis in B (FDR < 0.05, EdgeR). **F** Locus restricted differential analysis of *TMPRSS2*-linked peaks with increased accessibility in AT2 with aging (top panel in 3E). Black asterisk, p < 0.05 (T-test); Red asterisk, FDR < 0.05 (EdgeR) from dynamic peak analysis in B. **G** Annotation of motifs and evidence for transcription factor association within age-increased peaks. Blue bar, Motif present (FIMO); Green bar, Motif present (FIMO) and transcription factor association (ChIP-Atlas). **H** Overlap with ENCODE histone modification ChIP-seq data (Consortium, 2012) from SCREEN. **I** Genome browser representation of four *TMPRSS2*-linked peaks across age groups.

We identified two clusters of AT2 sites exhibiting increasing accessibility with age including several sites at candidate genes for SARS-CoV-2 host genes (cIII 30yo enriched and cIV 3yo + 30yo) (Figure 3B, Figure S6A, B). Intriguingly, these two clusters were enriched for processes related to viral infection and immune response such as viral release from host cell, interferon-gamma mediated signaling pathway, and positive regulation of ERBB signaling pathway (Figure 3C, Supplementary Table 6). Also, these age-dependent clusters were also enriched for phenotypes substantiated in mouse studies, such as pulmonary epithelial necrosis, increased monocyte cell number, and chronic inflammation (Fig. 3C, Supplementary Table 6). Further supporting an immune association with age-related chromatin accessibility in AT2 cells, we observed an enrichment of sequence motifs within these clusters for transcription factors involved in immune signaling such as STAT, IRF, and FOS/JUN (Figure 3D, Supplementary Table 7).

We focused on the *TMPRSS2* locus and determined how many of the 37 accessible chromatin sites co-accessible with the *TMPRSS2* promoter (in Figure 2D) showed increased accessibility with age in AT2 cells. We identified 13 sites with age-increased accessibility, of which 10 had significant effects (FDR < 0.05 via EdgeR and/or p < 0.05 via t-test) (Figure 3E, F, Figure S6, Supplementary Table 5). Age-increasing sites linked to *TMPRSS2* harbored sequence motifs for transcription factors such as NKX, FOXA, CEBPA, and inflammation-related factors such as STAT, IRF, and FOS/JUN (Figure 3G) many of which were corroborated by available ChIP-seq data in lung related samples (Oki et al., 2018). Furthermore, at 12 of the 13 age-increasing sites, we uncovered additional evidence for enhancer-related histone modifications from ENCODE supporting that they have *cis*-regulatory activity (Figure 3H) (Consortium, 2012). When viewed in genomic context these sites showed a clear age-dependent increase in read depth likely reflecting a higher fraction of accessible nuclei (Figure 3I).

### Genetic variants predicted to affect age-increased *TMPRSS2* sites are associated with respiratory phenotypes and *TMPRSS2* expression

Mapping the discrete accessible chromatin sites at genes required for SARS-CoV-2 viral entry allowed us to next characterize non-coding sequence variation that might affect regulation of these sites and contribute to phenotypic differences in the risk of lung disease. In particular, we focused on the 37 sites linked to *TMPRSS2* activity including 13 with age-increased chromatin accessibility.

In total, 8,002 non-singleton sequence variants in the gnomAD v3 database (Karczewski et al., 2019) overlapped a site either linked to or within 250kb of the *TMPRSS2* promoter. To determine which of these variants might affect regulatory activity in AT2 cells, we applied a machine learning approach (deltaSVM) (Lee et al., 2015) to model AT2 chromatin accessibility and predict variants with allelic effects on chromatin (see **Methods**). We identified 721 variants with significant effects (FDR<0.1) on AT2 chromatin accessibility, of which 148 mapped in an age-dependent site linked to *TMPRSS2* (Figure 4A). Among these 148 variants, 14 were common (defined here as minor allele frequency > 1%) in at least one major population group in gnomAD, several of which were predicted to disrupt AT2 age-dynamic TF motifs such as FOS/JUN, IRF, STAT, RUNX, NKX and ESR1 (Figure 4A). The common variants generally had consistent frequencies across populations, except for rs35074065 which was much less common in East Asians (EAS) relative to other populations (MAF=0.005, Figure 4B).

**Figure 4.**
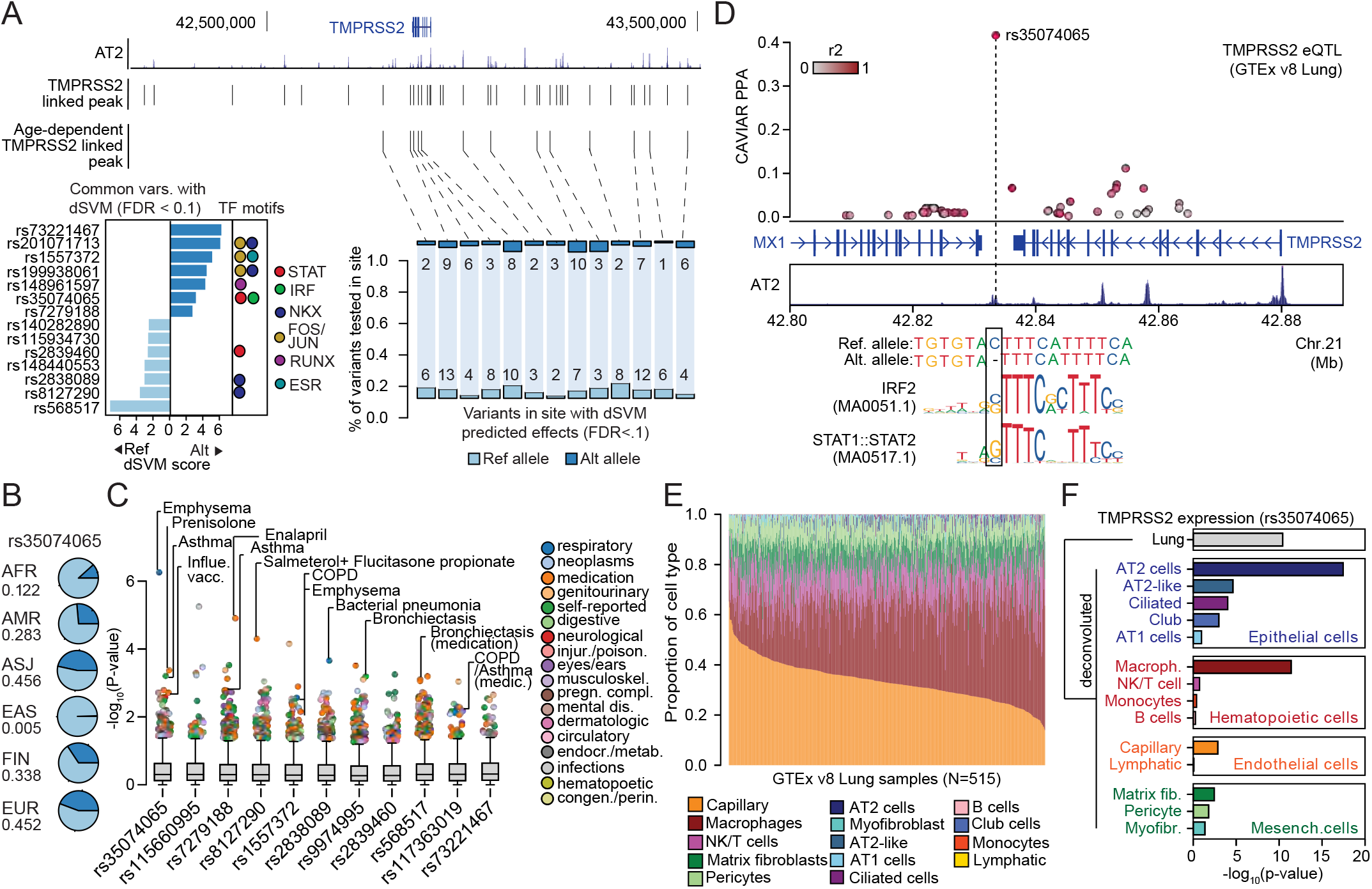
Genetic variants predicted to affect age-increasing AT2 accessible chromatin are associated with respiratory phenotypes and *TMPRSS2* expression. **A** Top: genome browser view of sites linked to *TMPRSS2* activity including those with age-dependent increase in activity. Right: Non-singleton genetic variants in gnomAD v3 mapping in each age-dependent site with predicted effects (FDR<.10) on AT2 chromatin accessibility using deltaSVM. Variants within each site are organized based on whether the reference (ref) or alternate (alt) allele has a higher predicted effect. Left: DeltaSVM scores of variants with predicted effects on AT2 chromatin accessibility and common (defined as MAF>1%) in at least one major population group in gnomAD v3, annotated with sequence motifs overlapping the variant for TF families enriched in age-increased AT2 sites. **B** Population frequency of variant rs35074065, which had predicted AT2 effects and was present at much lower frequency in East Asians relative to other population groups. AFR: African, AMR: Latino/American, ASJ: Ashkenazi Jewish, EAS: East Asian, FIN: Finnish, EUR: European (non-Finnish). **C** Association of common variants with predicted AT2 effects with human phenotypes in the UK Biobank. The majority of tested variants show at least nominal evidence (p<0.005) for association with phenotypes related to respiratory disease, infection and medication. **D** Fine-mapping probabilities for an *TMPRSS2* expression QTL in human lung samples from the GTEx project release v8. The variant rs35074065 has the highest casual probability (PPA=.42) for the eQTL, maps in an age-dynamic AT2 site and is predicted to disrupt binding of IRF and STAT TFs. Variants are colored based on r2 with rs35074065 in 1000 Genomes Project data using all populations. **E** Estimated cell type proportions for 515 human lung samples from GTEx derived using cell type-specific expression profiles for cell types with more than 500 cells from snRNA-seq data generated in this study. **F** Association p-values between rs35074065 genotype and *TMPRSS2* lung expression after including an interaction term between genotype and estimated cell type proportions for each sample. We observed stronger eQTL association when including an interaction with AT2 cell proportion as well as macrophage proportion.

We next determined whether common variants with predicted AT2 regulatory effects were associated with phenotypes related to respiratory function, infection, medication use or other traits using GWAS data generated using the UK Biobank (UKBB) (Sudlow et al., 2015). Across the 11 variants tested for association in UKBB data, the most significant association was between rs35074065 and emphysema (p=5.64×10^−7^) (Figure 4C). This variant was also more nominally associated (p<0.005) with asthma (p=6.7×10^−4^) and influenza vaccine (p=1.76×10^−3^). Furthermore, the majority of tested variants (8/11) were nominally associated (p<1×0^−3^) with at least one phenotype related to respiratory function or respiratory medication use including salmeterol + fluticasone propionate, which is commonly used to treat asthma and COPD (rs7279188 p=1.3×10^−5^), bacterial pneumonia (rs2838089 p=2.4×10^−4^), bronchiectasis (rs9974995 p=7.1×10^−4^, rs568517 p=8.1×10^−4^), and COPD (rs1557372 p=2.9×10^−3^) (Figure 4C).

Given that common AT2 variants showed predicted regulatory function and association with respiratory disease and infection phenotypes, we next asked whether these variants regulated the expression of *TMPRSS2* using human lung eQTL data from the GTEx v8 release. Among variants tested for association in GTEx, we observed a highly significant eQTL for *TMPRSS2* expression at rs35074065 (p=3.9×10^−11^) as well as more nominal eQTL evidence at rs1557372 (p=2.9×10^−5^) and rs9974995 (p=3.5×10^−6^). Furthermore, in fine-mapping data, rs35074065 had a high posterior probability (PPA=41.6%) and therefore likely has a direct casual effect on *TMPRSS2* expression (Figure 4D). This variant further disrupted sequence motifs for IRF and STAT transcription factors, suggesting that its effects may be mediated through interferon signaling and anti-viral programs (Figure 4D).

As the *TMPRSS2* eQTL at rs35074065 was identified in bulk lung samples, we finally sought to determine the specific cell types driving the effects of this eQTL. Using cell type-specific expression profiles derived from our snRNA-seq data, we estimated the proportions of 14 different cell types present in the 515 bulk lung RNA-seq samples from GTEx v8 (Figure 4E) (Aguet et al., 2019). We then tested the association between rs35074065 and *TMPRSS2* expression while including estimated cell type proportions for each sample in the eQTL model (see **Methods**). We observed highly significant association when including AT2 cell proportion (p=3.8×10^−18^) as well as macrophage proportion (p=4.0×10^−12^), supporting the possibility that the *TMPRSS2* eQTL at rs35074065 acts through AT2 cells and macrophages (Figure 4F).

## DISCUSSION

In this study, we focused on the lung, the organ at the center of COVID-19 morbidity and mortality. We generated a snATAC-seq reference dataset of the healthy human lung at three postnatal stages, and in parallel generated snRNA-seq data from the same samples to allow comparison with gene expression. Importantly, datasets were produced using uniform tissue procurement and single nucleus technologies for both modalities across samples. This consistency allowed us to uncover age-associated dynamics in gene expression and regulation. While we focus on COVID-19 related genes in this study, the datasets more broadly enable in-depth analysis of cell-type resolved dynamics of chromatin accessibility and gene expression in the human lung. We hope these datasets will be further utilized by the community to enhance knowledge and treatment of lung diseases.

One of the strongest findings that has been corroborated by multiple large-scale epidemiological studies is that infants and children, while still susceptible to infection, generally do not develop symptoms as severe as adults (Bi et al., 2020; CDC, 2020). Although the underlying molecular basis of this skew is unclear and is likely multifactorial, our data demonstrate that *ACE2*^+^ and *TMPRSS2^+^ and ACE2^+^/TMPRSS2*^+^ are detected in a higher proportion of AT2 nuclei in adult samples compared to the younger samples. These findings suggest that SARS-CoV-2 may enter proportionally fewer cells in younger lungs compared to adult lungs, leading to tempered viral replication and damage. While we await clinical validation of this finding, this difference in viral entry factors, in addition to likely differences in immune response to viral infection, may explain the age-related bias in COVID-19 severity.

The observed increase in the proportion of cells expressing viral entry genes is further corroborated by age-related changes in accessible chromatin, which offers insight for using gene regulatory mechanisms to restrict the expression of viral entry genes. For example, at the *TMPRSS2* locus we identified 10 accessible chromatin sites that showed significantly increased accessibility with age. These sites may therefore represent *cis* regulatory elements that contribute to activation of *TMPRSS2* gene expression in an increasing number of cells in adults and represent possible sites to modulate in order to restrict expression. Furthermore, one of the age-dependent sites harbors a sequence variant (rs35074065) significantly associated with *TMPRSS2* expression and respiratory phenotypes, suggesting it may be of particular value in this context.

To explore potential avenues for manipulating the expression of viral entry genes, we identified transcription factors enriched in sites with increased chromatin accessibility in adult AT2 cells compared to younger AT2 cells. These included transcription factors involved in stress and immune responses. For example, key interferon pathway-related factors STAT and IRF have binding sites in the 10 age-increased *TMPRSS2* peaks. The likely causal *TMPRSS2* eQTL variant rs35074065 is predicted to disrupt STAT and IRF binding, raising the possibility that STAT and/or IRF binding at this site may directly control *TMPRSS2* gene expression.

While our findings suggest that interferon pathway transcription factors may play a role in regulating the expression of SARS-CoV-2 entry genes such as *TMPRSS2*, extensive preclinical studies are needed to validate this regulation in an *in vivo* context. As a key anti-viral factor, interferon is stimulated in host cells upon infection by viruses, likely including SARS-CoV-2 (Lukhele et al., 2019; Mesev et al., 2019; Xia et al., 2018). The literature contains conflicting data regarding whether and how viral infection may act through the interferon pathway to regulate viral entry gene expression. For example, binding of the original SARS-CoV spike protein to ACE2 receptor in mice led to reduced *Ace2* expression in the lung (Kuba et al., 2005). However, a recent single-cell study suggested that viral-induced interferon activation stimulates *ACE2* expression (Ziegler, 2020). We caution that the potential effect of interferon signaling on COVID-19 needs to be investigated beyond viral entry, as the pathway likely has distinct roles in the different phases of the disease.

In our lung snRNA-seq data, *ACE2* is detected in a very small number of cells, a finding that is corroborated by a number of recent single cell studies (Qi et al., 2020; Waradon Sungnak, 2020; Zhao et al., 2020; Ziegler, 2020; Zou et al., 2020). The low fraction of nuclei that are *ACE2* positive could be due to low overall expression which in turn results in significant dropout in single cell or single nucleus RNA-seq. This suggests the possibility that ACE2 may not be needed at high levels for viral attachment to host cells. Alternatively, it is plausible that alternative receptors such as BSG also facilitate SARS-CoV-2 attachment *in vivo*. Compared to *ACE2, BSG* is expressed and co-expressed with proteases in a higher fraction of nuclei in AT2 and in additional cell types in the human lung.

To limit SARS-CoV-2 infection by manipulating the expression of viral entry proteins, we caution that inhibiting *ACE2* expression should not be a recommended strategy. Aside from being a viral receptor gene, *ACE2* is also required for protecting the lung from injury-induced acute respiratory distress phenotypes, the precise cause of COVID-19 mortality (Imai et al., 2005). Thus, inhibiting *ACE2* expression may compromise the ability of the lung to sustain damage. In comparison, *Tmprss2* mutant mice show no defects at baseline and are more resistant to the original SARS-CoV infection (Iwata-Yoshikawa et al., 2019; Kim et al., 2006). Thus, manipulating the expression of genes such as *TMPRSS2* may represent a safer path to limit SARS-CoV-2 viral entry. *TMPRSS2* is also involved in the entry of other respiratory viruses such as influenza, suggesting that modulating its expression may also be effective in deterring entry and spread of other viruses (Limburg et al., 2019).

In this study, we present the first snATAC-seq dataset of the human lung and complementary snRNA-seq data from the same samples. Here, we used COVID-19 genes to demonstrate how this dataset can be utilized. As COVID-19 GWAS data emerge, our datasets will offer a powerful cell type-resolved platform to interrogate mechanisms that may underlie genetic differences in the susceptibility and response to SARS-CoV-2 infection. Furthermore, our results suggest that modulation of the interferon pathway is a possible avenue to restrict *TMPRSS2* expression and viral entry. Identification of regulators that restrict the expression of viral entry genes without detrimentally affecting other aspects of the normal antiviral response will be a safe and effective strategy towards combating COVID-19. We note that this work is a product of the NHLBI-funded LungMap consortium, and our joint goal is to provide the community with fundamental knowledge of the human lung to help combat COVID-19.

## ACKNOWLEDGEMENTS

We are extremely grateful to the families who have generously given such precious gifts to support this research. We thank all the members of the LungMAP Consortium for their collaborations. We thank Dr. Bing Ren, Dr. Maike Sander, members of the Sun lab, Gaulton lab, Ren lab and the UCSD Center for Epigenomics for insightful discussions. We thank S. Kuan for sequencing and B. Li for bioinformatics support. We thank K. Jepsen and the UCSD IGM Genomics Center for sequencing the snRNAseq libraries. We thank the QB3 Macrolab at UC Berkeley for purification of the Tn5 transposase.

## AUTHOR CONTRIBUTIONS

Conceptualization: A.W., K.J.G., S.P., X.S.; Methodology: A.W., K.J.G., S.P., K.Z.; Validation: J.M.V., R.E.Y., J.B.; Formal analysis: A.W., K.J.G., S.P., J.C., O.B.P., J.B., M.J.V., J.M.N., M.G., Y.X.; Investigation: J.B., J.M.V., X.H.; Resources: R.M., H.H., L.R., C.P., G.P., J.A.W.; Data curation: A.W., K.J.G., S.P., J.C., O.B.P., J.B., M.J.V., P.K., D.A.F; Writing: A.W., J.C., J.M.V., K.J.G., S.P., X.S.; Supervision: A.W., K.J.G., S.P., J.A.W., X.S.; Project administration: A.W., K.J.G., S.P., J.M.V., X.S.; Funding acquisition: A.W., K.J.G., S.P., J.M.V., X.S.

## METHODS

### Human subjects and tissue collection

Donor lung samples were provided through the federal United Network of Organ Sharing via National Disease Research Interchange (NDRI) and International Institute for Advancement of Medicine (IIAM) and entered into the NHLBI LungMAP Biorepository for Investigations of Diseases of the Lung (BRINDL) at the University of Rochester Medical Center overseen by the IRB as RSRB00047606, as previously described (Ardini-Poleske et al., 2017; Bandyopadhyay et al., 2018). Portions (0.25-1.0 cm^3^) of small airway region of right middle lobe (RML) lung tissue were frozen in cryovials over liquid nitrogen and placed at −80°C for storage. Upon request, while kept frozen on dry ice, a tissue piece (approximately 100 mg) was chipped off the sample. These smaller samples were then shipped in cryovials to UCSD on an abundance of dry ice.

### Single nucleus ATAC-seq data generation

Combinatorial barcoding single nucleus ATAC-seq was performed as described previously with modifications (Cusanovich et al., 2015; Fang et al., 2019; Preissl et al., 2018) and using new sets of oligos for tagmentation and PCR (Supplementary Table 8). Briefly, for each sample, lung tissue was homogenized using mortar and pestle on liquid nitrogen. 1 ml nuclei permeabilization buffer (10mM Tris-HCL (pH 7.5), 10mM NaCl, 3mM MgCl2, 0.1% Tween-20 (Sigma), 0.1% IGEPAL-CA630 (Sigma) and 0.01% Digitonin (Promega) in water (Corces et al., 2017)) was added to 30 mg of ground lung tissue and tissue was resuspended by pipetting for 8-15 times. Nuclei suspension was incubated for 10 min at 4°C and filtered with 30 μm filter (CellTrics). Nuclei were pelleted with a swinging bucket centrifuge (500 x g, 5 min, 4°C; 5920R, Eppendorf) and resuspended in 500 μL high salt tagmentation buffer (36.3 mM Tris-acetate (pH = 7.8), 72.6 mM potassium-acetate, 11 mM Mg-acetate, 17.6% DMF) and counted using a hemocytometer. Concentration was adjusted to 2,000 nuclei/9 μl, and 2,000 nuclei were dispensed into each well of one 96-well plate. For tagmentation, 1 μL barcoded Tn5 transposomes (Fang et al., 2019) was added using a BenchSmart™ 96 (Mettler Toledo), mixed five times and incubated for 60 min at 37 °C with shaking (500 rpm). To inhibit the Tn5 reaction, 10 μL of 40 mM EDTA were added to each well with a BenchSmart™ 96 (Mettler Toledo) and the plate was incubated at 37 °C for 15 min with shaking (500 rpm). Next, 20 μL 2 x sort buffer (2 % BSA, 2 mM EDTA in PBS) was added using a BenchSmart™ 96 (Mettler Toledo). All wells were combined into a FACS tube and stained with 3 μM Draq7 (Cell Signaling). Using a SH800 (Sony), 20 2n nuclei were sorted per well into eight 96-well plates (total of 768 wells) containing 10.5 μL EB (25 pmol primer i7, 25 pmol primer i5, 200 ng BSA (Sigma). Preparation of sort plates and all downstream pipetting steps were performed on a Biomek i7 Automated Workstation (Beckman Coulter). After addition of 1 μL 0.2% SDS, samples were incubated at 55 °C for 7 min with shaking (500 rpm). 1 μL 12.5% Triton-X was added to each well to quench the SDS. Next, 12.5 μL NEBNext High-Fidelity 2× PCR Master Mix (NEB) were added and samples were PCR-amplified (72 °C 5 min, 98 °C 30 s, (98 °C 10 s, 63 °C 30 s, 72°C 60 s) × 12 cycles, held at 12 °C). After PCR, all wells were combined. Libraries were purified according to the MinElute PCR Purification Kit manual (Qiagen) using a vacuum manifold (QIAvac 24 plus, Qiagen) and size selection was performed with SPRI Beads (Beckmann Coulter, 0.55x and 1.5x). Libraries were purified one more time with SPRI Beads (Beckmann Coulter, 1.5x). Libraries were quantified using a Qubit fluorimeter (Life technologies) and the nucleosomal pattern was verified using a Tapestation (High Sensitivity D1000, Agilent). The library was sequenced on a HiSeq4000 or NextSeq500 sequencer (Illumina) using custom sequencing primers with following read lengths: 50 + 10 + 12 + 50 (Read1 + Index1 + Index2 + Read2). Primer and index sequences are listed in Supplementary Table 8.

### Single nucleus RNA-seq data generation

Droplet-based Chromium Single Cell 3’ solution (10x Genomics, v3 chemistry)(Zheng et al., 2017) was used to generate snRNA-seq libraries. Briefly, 30 mg pulverized lung tissue was resuspended in 500 μl of nuclei permeabilization buffer (0.1% Triton X-100 (Sigma-Aldrich, T8787), 1X protease inhibitor, 1 mM DTT, and 0.2 U/μl RNase inhibitor (Promega, N211B), 2% BSA (Sigma-Aldrich, SRE0036) in PBS). Sample was incubated on a rotator for 5 minutes at 4°C and then centrifuged at 500 rcf for 5 minutes (4°C, run speed 3/3). Supernatant was removed and pellet was resuspended in 400 μl of sort buffer (1 mM EDTA 0.2 U/μl RNase inhibitor (Promega, N211B), 2% BSA (Sigma-Aldrich, SRE0036) in PBS) and stained with DRAQ7 (1:100; Cell Signaling, 7406). 75,000 nuclei were sorted using a SH800 sorter (Sony) into 50 μl of collection buffer consisting of 1 U/μl RNase inhibitor in 5% BSA; the FACS gating strategy sorted based on particle size and DRAQ7 fluorescence. Sorted nuclei were then centrifuged at 1000 rcf for 15 minutes (4°C, run speed 3/3) and supernatant was removed. Nuclei were resuspended in 35 μl of reaction buffer (0.2 U/μl RNase inhibitor (Promega, N211B), 2% BSA (Sigma-Aldrich, SRE0036) in PBS) and counted on a hemocytometer. 12,000 nuclei were loaded onto a Chromium controller (10x Genomics). Libraries were generated using the Chromium Single Cell 3’ Library Construction Kit v3 (10x Genomics, 1000078) according to manufacturer specifications. CDNA was amplified for 12 PCR cycles. SPRISelect reagent (Beckman Coulter) was sued for size selection and clean-up steps. Final library concentration was assessed by Qubit dsDNA HS Assay Kit (Thermo-Fischer Scientific) and fragment size was checked using Tapestation High Sensitivity D1000 (Agilent) to ensure that fragment sizes were distributed normally about 500 bp. Libraries were sequenced using the NextSeq500 and a HiSeq4000 (Illumina) with these read lengths: 28 + 8 + 91 (Read1 + Index1 + Read2).

### Single nucleus RNA-seq analysis

Sequencing reads were demultiplexed (cellranger mkfastq) and processed (cellranger count) using the Cell Ranger software package v3.0.2 (10x Genomics). Reads were aligned to the human reference hg38 (Cell Ranger software package v3.0.2). Reads mapping to intronic and exon sequences were retained. Resulting UMI feature-barcode count matrices were loaded into Seurat (Stuart et al., 2019). All genes represented in >=3 nuclei and cells with 500-4000 detected genes were included for downstream processing. UMI counts were log-normalized and scaled by a factor of 10,000 using the NormalizeData function. Top 3000 variable features were identified using the FindVariableFeatures function and finally scaled using the ScaleData function. Barcode collisions were removed for individual datasets using DoubletFinder (McGinnis et al., 2019) with following parameters: pN =0.15 and pK = 0.005, anticipated collision rate = 10%. Clusters were assigned a doublet score (pANN) and classification as “doublet” or “singlet”; called doublets and cells with a pANN score > 0 were removed. UMI matrices for datasets were merged and corrected for batch effects due to experiment date, donor, and sex using the Harmony package (Korsunsky et al., 2019). UMAP coordinates and clustering were performed using the RunUMAP, FindNeighbors, and FindClusters functions in Seurat with principal components 1-23. 25-26, and 28. Clusters were annotated, and putative doublets as defined by expression of canonically mutually exclusive markers were excluded from analysis; remaining cells were re-clustered using the previously described parameters. Final cluster annotation was done using canonical markers. For genes of interest such as (e.g. *ACE2*, *TMPRSS2*), nuclei with at least one UMI for the gene were considered “expressing”. To analyze changes in percentage of nuclei expressing we performed One-way ANOVA (ANalysis Of VAriance) with post-hoc Tukey HSD (Honestly Significant Difference) using GraphPad Prism version 8.0.0 for Windows, GraphPad Software, San Diego, California USA, www.graphpad.com. Due to one potential outlier in the 30wkGA group (D062) we performed in addition a simple t-test comparing 3 yr to 30 yr groups. Differential gene expression analysis between *ACE2*^+^ and *ACE2*^-^ AT2 cells we used FindAllMarkers with parameters logfc = 0, min.pct = 0, test.use = “wilcox”, verbose = TRUE.

### Normalization and comparison of gene expression frequency across snRNA-seq and scRNA-seq datasets

Single cell RNA-seq (10x Genomics 3’ v2) of 4 aged (>55yr) control lungs were obtained from publicly available data (Morse et al., 2019; Reyfman et al., 2019). Raw gene expression matrices were downloaded from Gene Expression Omnibus (GEO) repository (GSE128033 and GSE122960). Cells were filtered using the following commonly used criteria: >500 expressed genes and <10% UMIs mapped to mitochondrial DNAs. In addition, cells with greater than or equal to 40,000 UMIs were excluded from the downstream analysis; this filtration criterion was selected based on the distribution of UMIs in single cells in individual donors. Seurat (version 3) (Stuart et al., 2019) was used to identify AT2 cells from individual aged donors. Nuclei from the 9 libraries generated in this study and cells from libraries for the 4 aged donors were integrated using the Seurat 3 standard integration pipeline (Stuart et al., 2019).

We calculated *ACE2* and *TMPRSS2* expression frequency in AT2 cells (percentage of AT2 cells with >0 UMI) in individual donors. We then performed median based normalization, so all donors reached the same median value. In calculating the median value for each donor, the expression frequency values of genes (n=26,260) common in both datasets were used.

### Single nucleus ATAC-seq analysis

For each sequenced snATAC-Seq libraries, we obtained four FASTQ files paired-end DNA reads as well as the combinatorial indexes for i5 (768 different PCR indices) and T7 (96 different tagmentation indices; Supplementary Table 8). We selected all reads with <= 2 mistakes per individual index (Hamming distance between each pair of indices is 4) and subsequently integrated the full barcode at the beginning of the read name in the FASTQ files (https://gitlab.com/Grouumf/ATACdemultiplex/). Next, we used trim galore (v.0.4.4) to remove adapter sequences from reads prior to read alignment. We aligned reads to the hg19 reference genome using bwa mem (v.0.7.17) (Li and Durbin, 2009) and subsequently used samtools (Li et al., 2009) to remove unmapped, low map quality (MAPQ<30), secondary, and mitochondrial reads. We then removed duplicate reads on a per-cell basis using MarkDuplicates (BARCODE_TAG) from the picard toolkit. As an initial quality cutoff, we set a minimum of 1,000 reads (unique, non-mitochondrial) and observed 120,090 cells passing this threshold.

We used a previously described pipeline to identify snATAC-seq clusters (Chiou et al., 2019). Briefly, we used scanpy (Wolf et al., 2018) to uniform read depth-normalize and log-transform read counts within 5 kb windows. We then identified highly variable (*hv*) windows (min_mean=0.01, min_disp=0.25) and regressed out the total read depth across *hv* windows (usable counts) within each experiment. We then merged cells across experiments and extracted the top 50 PCs, using Harmony (Korsunsky et al., 2019) to correct for potential confounding factors including donor-of-origin and biological sex. We used Harmony-corrected components to build a nearest neighbor graph (n_neighbors=30) using the cosine metric, which was used for UMAP visualization (min_dist=0.3) and Leiden clustering (resolution=1.5) (Traag et al., 2019).

Prior to the final clustering results, we performed iterative clustering to identify and remove cells mapping to clusters with aberrant quality metrics. First, we removed 3,183 cells mapping in clusters with low read depth. Next, we removed 20,718 cells mapping in clusters with low fraction of reads in peaks. Finally, we re-clustered the cells at high resolution and removed 5,209 cells mapping in potential doublet sub-clusters. On average, these sub-clusters had higher usable counts, promoter usage, and accessibility at more than one marker gene promoter. After removing all of these cells, our final clusters consisted of 90,980 cells. To identify marker genes for each cluster, we used linear regression models with gene accessibility as a function of cluster assignment and usable counts across single cells.

### Computing relative accessibility scores

We define an accessible locus as the minimal genomic region that can be bound and cut by the enzyme. We use *L* ⊂ *N* to represent the set of all accessible loci. We further define a pseudo-locus as the set of accessible loci that relates to each other in a certain meaningful way (for example, nearby loci, loci from different alleles). In this example, pseudo-loci correspond to peaks. We use {*d_i_* | *d_i_* ⊂ *L*} to represent the set of all pseudo-loci. Let *a_l_* be the accessibility of accessible locus *l*, where *l* ∈ *L*. We define the accessibility of pseudo-locus *d_i_* as *A_i_* = ∑_*k*∈*d_i_*_ *a_k_*, *i.e*., the sum of accessibility of accessible loci associated with di. Let *C_j_* be the library complexity (the number of distinct molecules in the library) of cell *j*. Assuming unbiased PCR amplification, then the probability of being sequenced for any fragment in the library is: 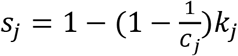, where *k_j_* is the total number of reads for cell *j*. If we assume that the probability of a fragment present in the library is proportional to its accessibility and the complexity of the library, then we can deduce that the probability of a given locus *l* in cell *j* being sequenced is: *p_lj_* ∝ *a_l_C_j_s_j_*. For any pseudo-locus *d_i_*, the number of reads in *d_i_* for cell *j* follows the Poisson binomial distribution, and its mean is 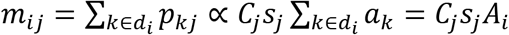. Given a pseudo-locus (or peak) by cell count matrix *O*, we have: 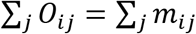. Therefore, 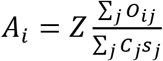, where *Z* is a normalization constant. When comparing across different samples the relative accessibility may be desirable as they sum up to a constant, *i.e*., ∑_*i*_ *A_i_* = 1 × 10^6^. In this case, we can derive 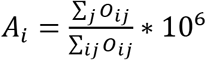.

### Calculating the relative percent of cells with accessibility at a locus

To correct for biases occurring from differential read depths between clusters, we used the following strategy to determine the relative ratio of cells with accessibility at a given locus. We defined the set of accessible loci *L* of a given dataset *D* as the genomic regions covered by the set peaks *P* inferred from *D*. We define *X* the set of cells from *D*, and *S* a partitioning of *X*. For a given partition *S_i_* ∈ *S* and for each feature *p_j_* ∈ *P*, we computed *m_ij_* the ratio of cells from *S_i_* with at least one read overlapping *p_j_*. We then defined the score *s_ij_*, of loci *p_j_* in *S_i_* as 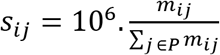. We finally define the relative ratio of cells normalized across the different clusters as 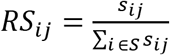.

### Associating promoters to candidate distal regulatory elements

To identify AT2 co-accessible loci with the promoters of *TMPRSS2, ACE2, FURIN, BSG*, and *CTSL* we used Cicero (Pliner et al., 2018). First, we performed a Cicero analysis for each individual cluster using a genomic window of 1 Mb (co-accessibility score >0.05). In addition, we performed Cicero using a random subset of 15,000 nuclei from the complete dataset and a genomic window of 250 kb (co-accessibility score >0.05). We then defined the promoter regions of *ACE2, TMPRSS2, FURIN, BSG*, and *CTSL* as transcriptional start site (TSS) +/- 1 kb and selected the sites co-accessible with each of the promoters (co-accessibility score >0.05). Finally, we merged the elements co-accessible with the gene promoters from both analyses to generate a union set of candidate elements.

### Identification and clustering of AT2 peaks with changes in chromatin accessibility genome-wide

We used edgeR (Robinson et al., 2010) to identify differential accessible peaks between each of pair of time points. As input we used the 122,352 peaks in AT2 cell. Dataset ID and sex were used as technical covariates. Sites with False Discovery Rate (FDR) < 0.05 after Benjamini-Hochberg correction were considered significant. Next, we performed K-means using the relative accessibility score with a *loci x timepoints* matrix. We used K from 5 to 8 and computed the Davis-Bouldin index to determine the best K to partition the loci. let 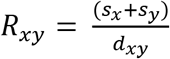 with *s_x_* the average distance of each sample from cluster *x* and *d_xy_* the distance between the centroids of clusters *x* and *y*. The Davies-Bouldin index is defined as 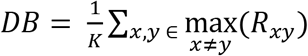 and low *DB* scores indicate better partitioning. We obtained an optimal partition with K=5.

### Identification of AT2 peaks with changes in chromatin accessibility at candidate gene loci

The ensemble of cells *X* from *D* can be divided per timepoint, cell subtype, or donor. We identified for individual donors the relative % of cells with at least one read in peaks associated with *ACE2, TMPRSS2, FURIN, BSG*, and *CTSL* promoters. As a background to calculate the relative % of cells, we used the merged set of peaks from all the clusters. Then, we computed a Student test for two independent samples with equal variance for each pair of categories: 30 wk^GA^, 3 yo and 30 yo. For each element the relative % of cells were used as measurement variable and the timepoint as nominal variable.

### Annotation of genomic elements

The GREAT algorithm (McLean et al., 2010) was used to annotate distal genomic elements using the following settings: 2 nearest gene within 1Mb.

### Transcription factor related analyses

*De novo* motif enrichment analysis in genomic elements was performed using HOMER (Heinz et al., 2010) with standard parameters. Motif scanning was performed using FIMO (Grant et al., 2011) online interface and default parameters. Motif files were downloaded from JASPAR (Fornes et al., 2020) in MEME format. Motifs scanned were MA0102.4 (CEBPA), MA0673.1(NKX2-8), MA0153.1(HNF1B), MA0503.1(NKX2-5), MA0877.2(BARHL1), PB0022.1(GATA5), MA0490.1(JUNB), PH0171.1(NKX2-1), MA0148.1(FOXA1), MA0144.1(STAT3), MA0517.1(STAT1::STAT2), MA0050.1(IRF), MA0007.2(AR), and MA0592.1(ESRRA). To identify overlap with TF ChIP-seq sites, we used ChIP-atlas (Oki et al., 2018). We downloaded a BED file for “TFs and other” antigens across all lung related samples from the Peak Browser. We intersected these peaks with the *TMPRSS2*-linked peaks and the FIMO motifs (Grant et al., 2011). In addition, we downloaded enhancer related histone modifications (H3K4me1, H3K27ac) from the SCREEN database and intersected with the peak lists (Consortium, 2012).

### Predicting variant effects on chromatin accessibility

We used deltaSVM (Lee et al., 2015) to predict the effects of variants on chromatin accessibility in AT2 cells. First, we extracted the sequences underlying AT2 sites that were promoter-distal (>±500 bp from GENCODE v19 transcript TSS for protein-coding and long non-coding RNA genes). As described previously (Chiou et al., 2019), we trained an AT2 sequence-based model and used it to predict effects for all possible combinations of 11mers. Next, to compile a comprehensive set of variants to test, we downloaded lists of variants from gnomAD v3 (Karczewski et al., 2019) and filtered out variants that were singletons or indels longer than 3 bp. We then used the liftOver (Tyner et al., 2017) utility to transform GRCh38 into GRCh37/hg19 coordinates. We retained variants from either dataset that mapped within *TMPRSS2* linked sites and extracted sequences in a 19 bp window around each variant (±9 bp flanking each side). Finally, we calculated deltaSVM z-scores for each variant by predicting deltaSVM scores, randomly permuting 11mer effects and re-predicting deltaSVM scores, and using the parameters of the null distribution to calculate deltaSVM z-scores. From the z-scores, we calculated p-values and q-values and defined variants with significant effects using a threshold of FDR<0.1. We identified common variants defined as minor allele frequency >.01 in at least one major population group. For each common variant, we obtained sequence surrounding each variant allele and predicted sequence motifs from the JASPAR database (Fornes et al., 2020) using FIMO (Grant et al., 2011), and focused on motifs of TF families enriched in age-dependent AT2 chromatin.

### Phenotype associations for predicted effect variants

We downloaded UK biobank round 2 GWAS combined sex results (Lab, 2020; Sudlow et al., 2015). We used broad disease categories from the ICD-10-CM to classify ICD10 phenotypes, except for ICD10 codes relating to unclassified symptoms, external causes of morbidity, and factors influencing health status and contact with health services. We combined all non-cancer, self-reported diseases into a single category (self-reported) as well as all treatments and medications (medication). We then extracted GWAS association results for variants that were not tagged as low confidence variants, had significant deltaSVM effects, and mapped in *TMPRSS2*-linked aging-related sites. From these variants, we removed one (rs199938061) which was in perfect linkage disequilibrium with another variant.

### Deconvoluting the *TMPRSS2* lung eQTL

We used MuSiC (v.0.1.1) (Wang et al., 2019) to estimate the proportions of lung cell types with >500 cells from our scRNA-seq dataset in lung bulk RNA-seq samples from the GTEx v8 release (Aguet et al., 2019). We combined cell type labels for capillary (distal and proximal), macrophages (M1 and M2), matrix fibroblasts (1 and 2), and NK/T cells. We modeled the relationship between TMM-normalized *TMPRSS2* expression as a function of the interaction between genotype and cell type proportion, while considering the covariates used in the original GTEx data including sex, sequencing platform, PCR, 5 genotype PCs, and 59 inferred PCs from the expression data. From the original inferred PCs, we excluded inferred PC 1 because it was highly correlated with AT2 cell type proportion (Spearman ρ=0.67).

## SUPPLEMENTARY FIGURE LEGENDS

**Figure S1. Quality control of snRNA-seq and snATAC-seq datasets. A** Representative UMI barcode distribution output from CellRanger pipeline for snRNA-seq libraries from human lung. **B** Number of nuclei passing quality control filtering for snRNA-seq libraries. **C** Genes detected per nucleus. **D** Sequencing saturation of snRNA-seq libraries. **E** Nuclei with less than 1,000 uniquely mapped reads were filtered from snATAC-seq datasets. **F** Number of nuclei passing quality control filtering for snATAC-seq libraries. **G** Average number of reads per nucleus. **H** Fraction of reads in peak regions per dataset. All data are represented as mean ± SD.

**Figure S2. Marker plots for cluster annotation and expression profiling of candidate genes involved in SARS-CoV-2 cell entry. A** Dot plot of marker genes used for cluster annotation. **B-D** Cell type specific gene expression of candidate genes for cell entry. Violin plots display expression values per nucleus for genes encoding **B** Cathepsin L (*CTSL*), **C** FURIN (*FURIN*) and **D** Basigin (BSG, CD147). **E** Correlation of *ACE2+* and *TMPRSS2+* AT2 cells with linear regression. **F, G** Fraction of AT2 cells with expression of *ACE2* and *TMPRSS2* at each time point. Data are the same as Fig. 1D, E, but with potential outlier sample D062 removed. * p <0.05 (One-way ANOVA with post-hoc Tukey test).

**Figure S3. Expression analysis of viral entry genes**. Displayed are violin plots of expression levels for entry genes related to other viruses including SARS-CoV, MERS, coronavirus associated with common cold, Rhinovirus, Respiratory Syncytial Virus (RSV), Adenovirus, Influenza Virus.

**Figure S4. Integrative analysis of *ACE2* and *TMPRSS2* expression in lungs from aged individuals. A** Seurat3 Standard Integration (Stuart et al., 2019) was applied to snRNA-seq data for 9 donors generated as part of this study and publicly available scRNA-seq datasets 4 additional donor lungs (age> 55). AT2 cells from 13 donors were clustered together via Louvain clustering with minimal batch variation. Left panel: t-SNE visualization of cells colored by major cell type annotation. Epi other: predicted non-AT2 epithelial cells. Right panel: t-SNE visualization of cells colored by donor information. **B** Normalized expression frequency of *ACE2* (left) and *TMPRSS2* (right) in AT2 cells. p value was calculated using one-tailed t-test comparing normalized frequency in donors of 30yo group and aged group.

**Figure S5. Marker plots for cluster annotation of snATAC-seq and profiling of peaks at candidate genes for SARS-CoV-2 cell entry. A** Dot plot of marker genes used for cluster annotation. **B-D** Cell type resolved chromatin accessibility at peaks within +/- 650 kb of candidate genes for cell entry. Displayed are data for **B** FURIN (*FURIN*) and **C** Basigin (*BSG*, CD147) **D** Cathepsin L (*CTSL*). Values are displayed as row normalized proportion of cells with a fragment in a peak region. Black asterisks denote co-accessibility from Cicero >0.05 (Cusanovich et al., 2018).

**Figure S6. Quantification of peaks with increased accessibility with age at tested loci and donor resolved activity of sites not increased at *TMPRSS2* locus. A** Number of peaks within +/- 650 kb of candidate genes for cell entry overlapping cIII and cIV from Figure 3B. **B** Number of peaks co-accessible with the promoter of candidate genes for cell entry overlapping cIII and cIV from Figure 3B. **C** Donor resolved analysis of 24/37 peaks at the *TMPRSS2* gene locus. Red asterisks denote FDR <0.05 (EdgeR) and black asterisks denote p < 0.05 via t-test.

## SUPPLEMENTARY TABLE LEGENDS

**Supplementary Table 1. Donor metadata tables**. Sheet 1: 30wk^GA^ - 30yo: Donor ID, age, sex, race, clinical pathology diagnosis (clinPathDx), gestational age, overall quality of the lung tissue assessment, type of death and cause of death were listed. Not shown are data on body weight, body height, total lung weight and radial alveolar count assessment of alveolarization. All were all within normal limits for age. Abbreviations: DCD: donor after cardiac death; DBD: donor after brain death; GA: gestational age; RDS: respiratory distress syndrome. Sheet 2: aged cohort: Donor ID, age, sex, smoking history, race and cause of death were listed (Morse et al., 2019; Reyfman et al., 2019).

**Supplementary Table 2**. Cluster composition and number and fraction of nuclei expressing candidate for SARS-CoV2 cell entry.

**Supplementary Table 3**. Differential expressed analysis between *ACE2*^+^ and *ACE2*^-^ as well as *TMPRSS2*^+^ and *TMPRSS2*^-^ AT2 cells.

**Supplementary Table 4**. Annotation of peaks within a window of +/- 650 kb of candidate genes for SARS-CoV2 cell entry.

**Supplementary Table 5**. Annotation of peaks co-accessible with candidate genes for SARS-CoV2 cell entry and age-associated changes of chromatin accessibility of peaks co-accessible with *TMPRSS2* promoter.

**Supplementary Table 6**. GREAT analysis of peaks increasing with age in AT2 cells (groups cIII and cIV in Fig 3B).

**Supplementary Table 7**. *De novo* motif enrichment analysis of peaks increasing with age in AT2 cells (groups cIII and cIV in Fig 3B).

**Supplementary Table 8**. Indexes and primer sequences for snATAC-seq libraries.

