## Supplementary Figures for "Single Nucleus Multiomic Profiling Reveals Age-Dynamic Regulation of Host Genes Associated with SARS-CoV-2 Infection"

### Supplementary Figure 1

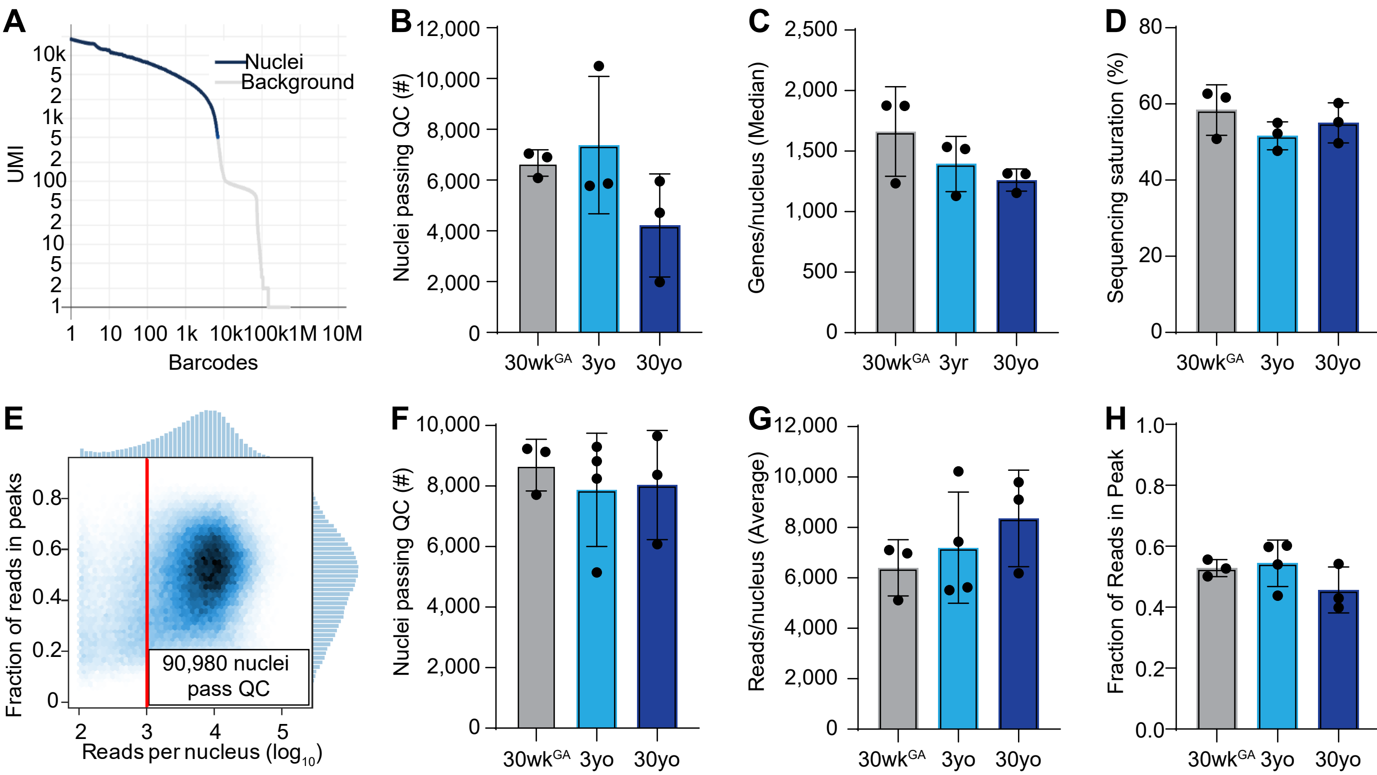

#### Supplementary Figure 2

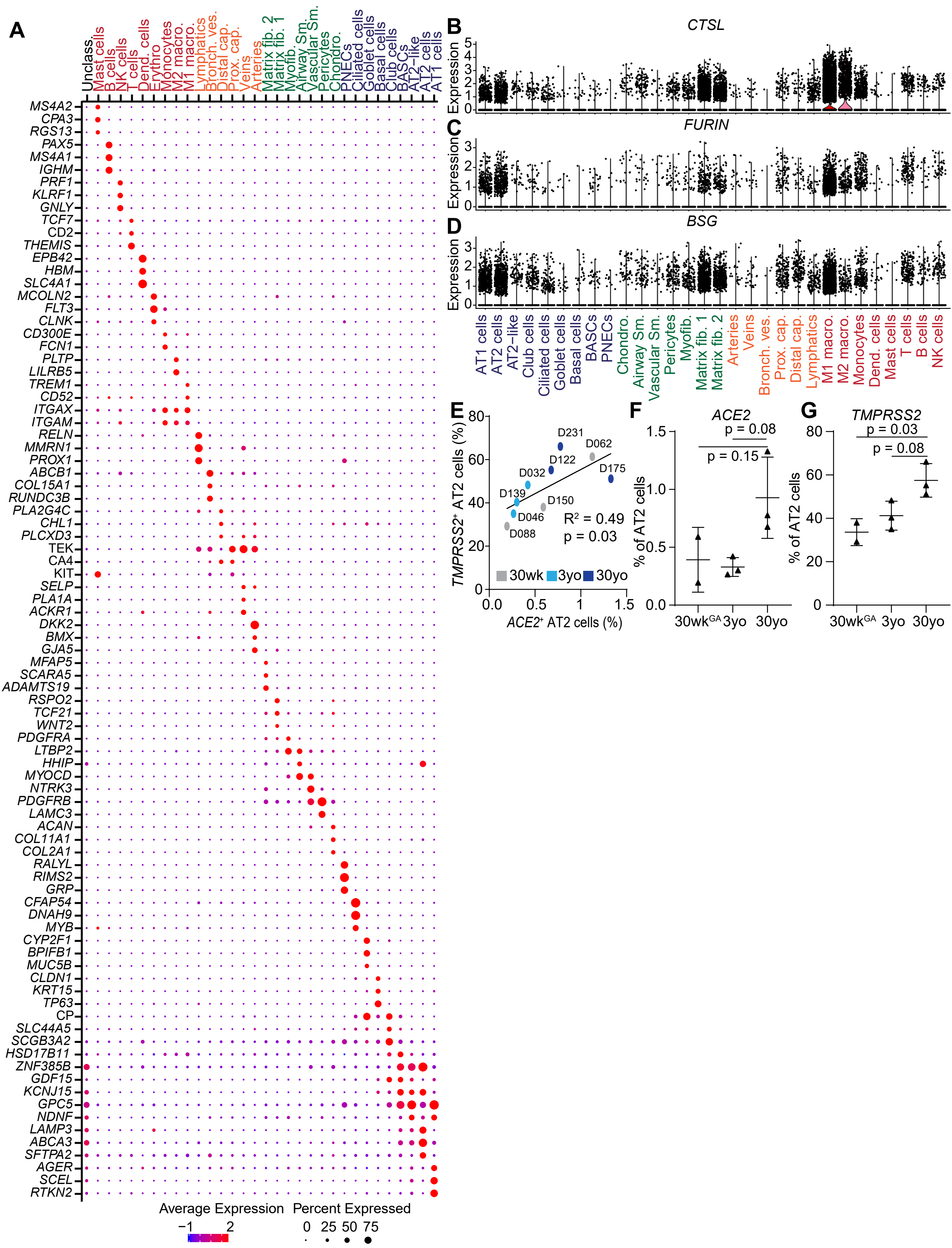

### Supplementary Figure 3

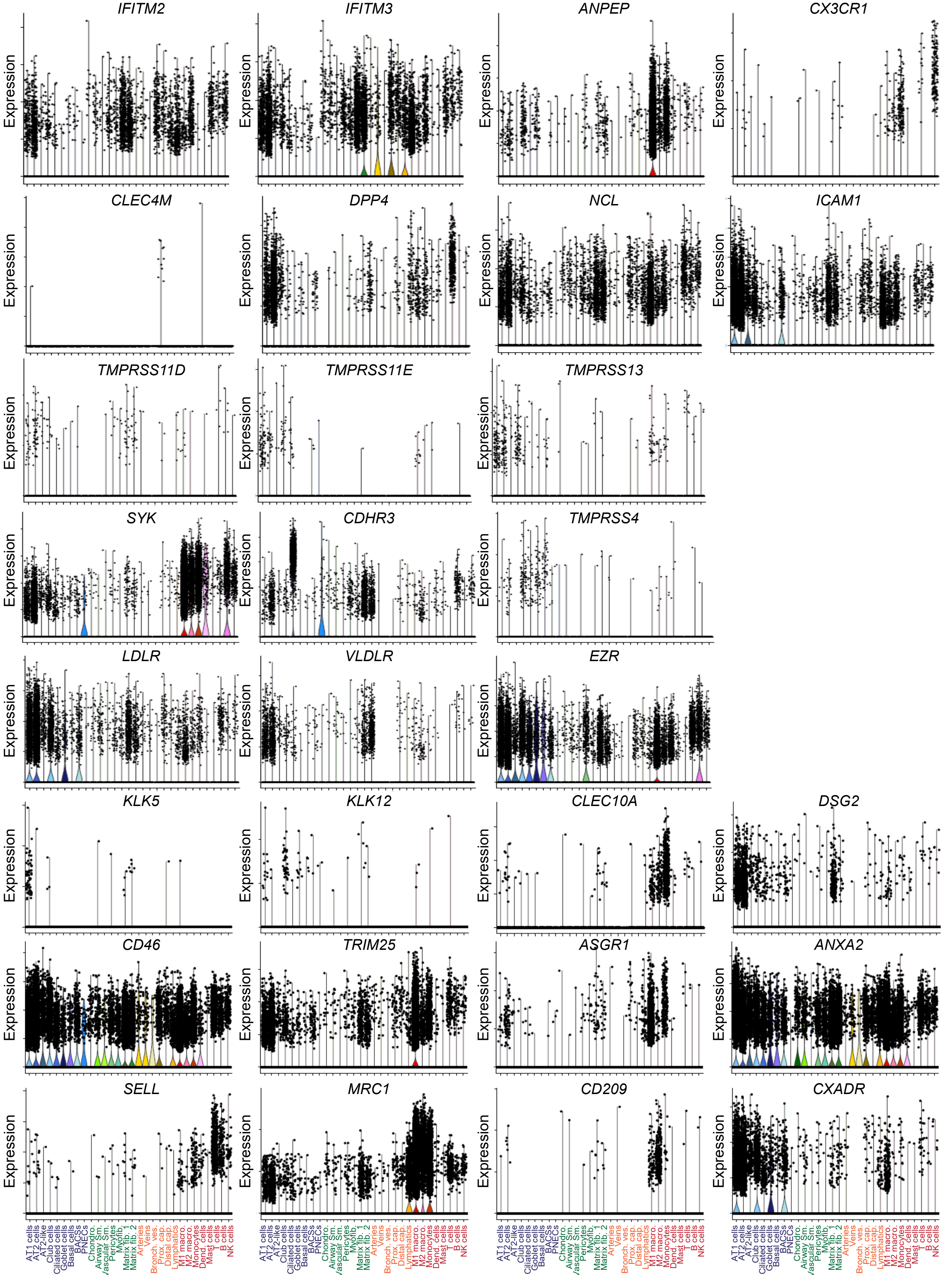

#### Supplementary Figure 4

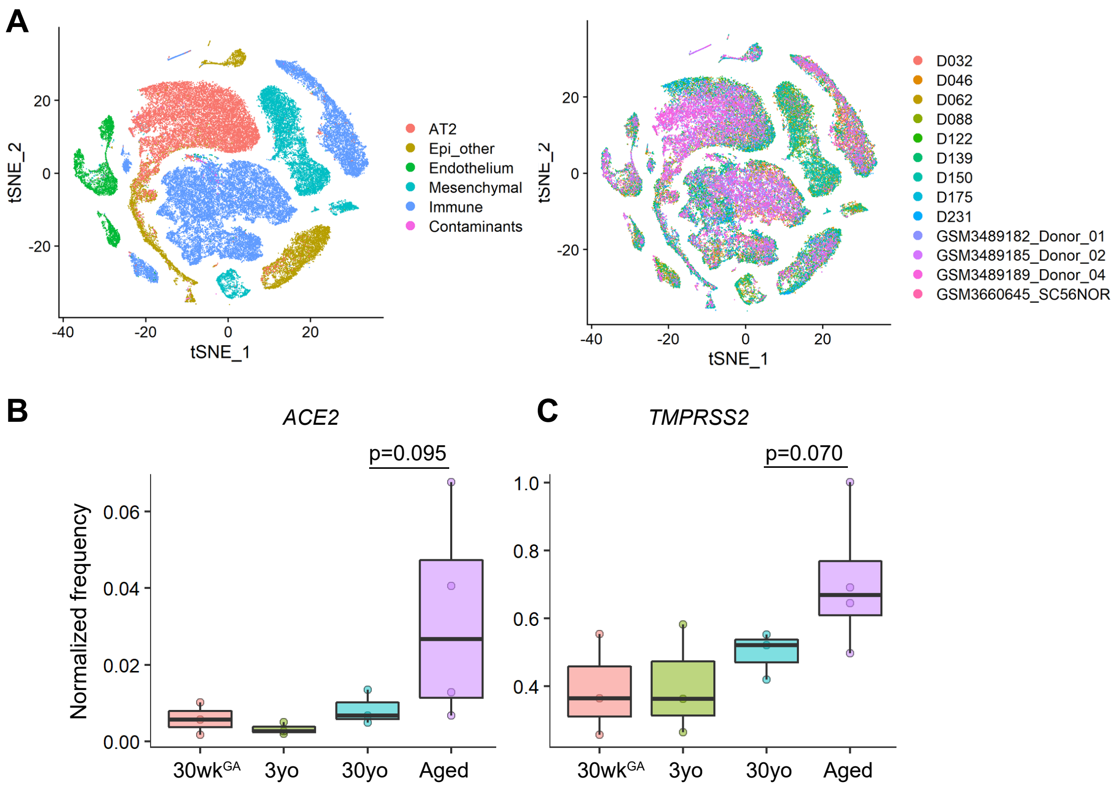

**Supplementary Figure 5**

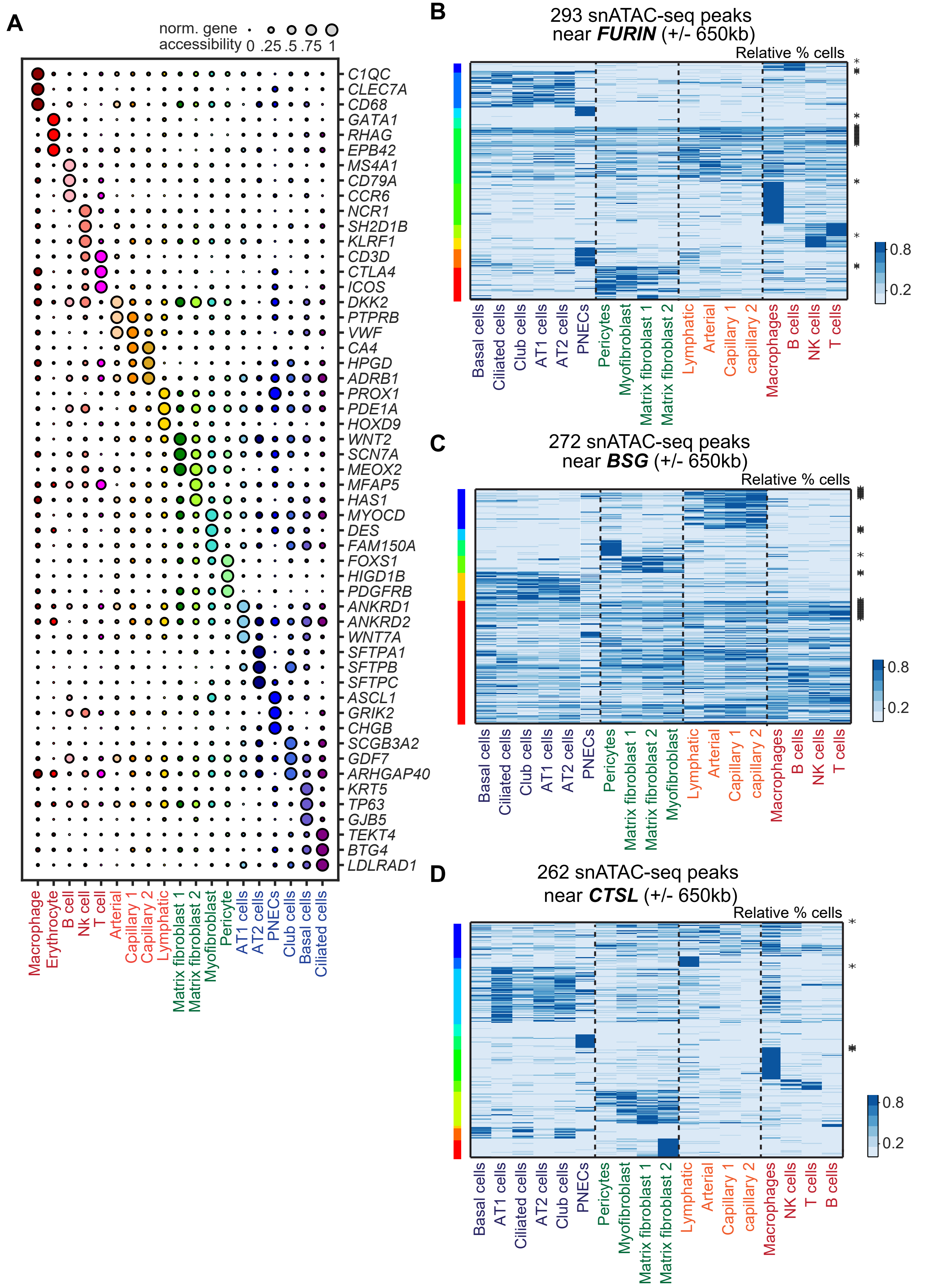

### Supplementary Figure 6

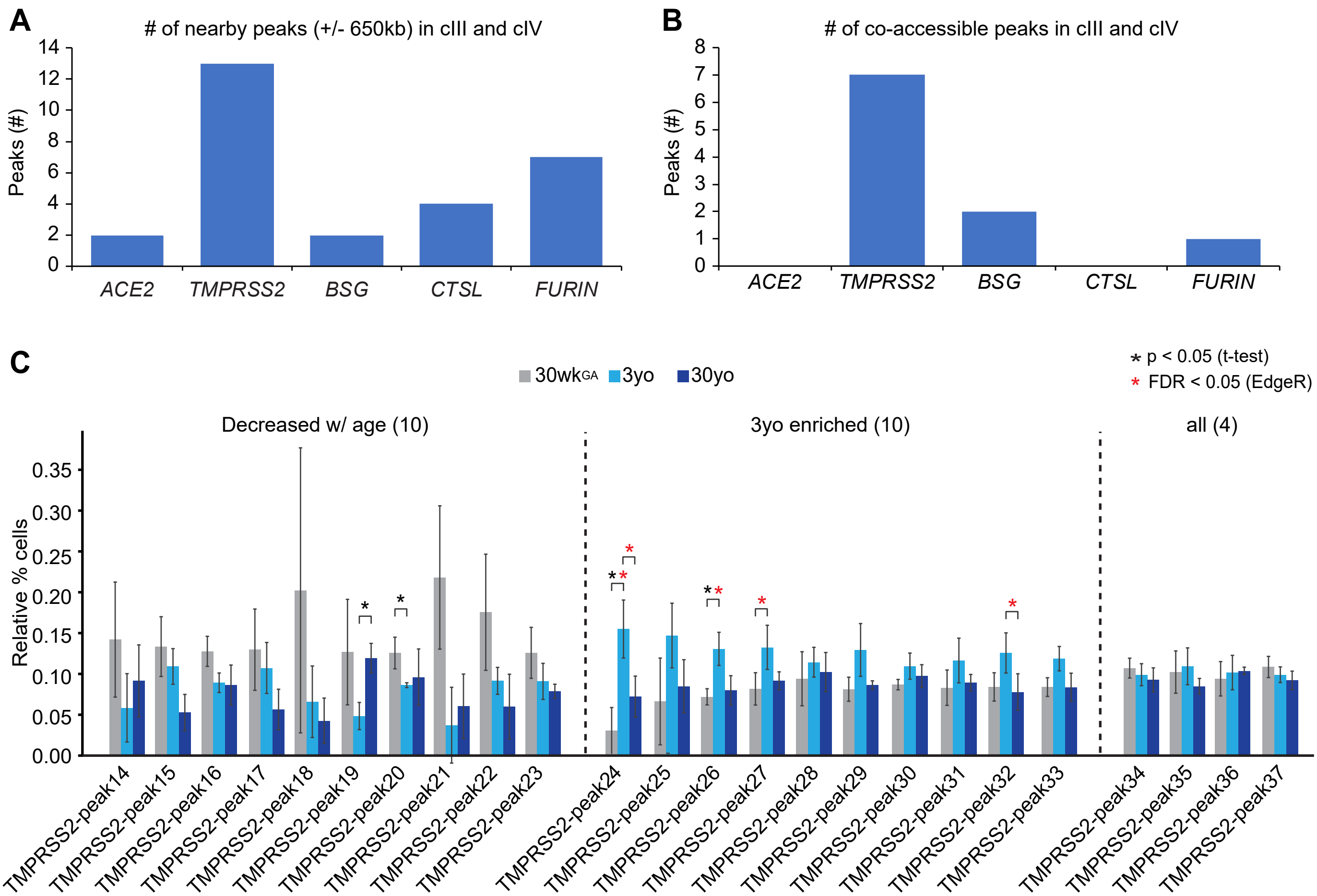
